# Emergence of diverse ligase reactivities from a single RNA evolution experiment

**DOI:** 10.64898/2026.01.15.699813

**Authors:** Annyesha Biswas, Saurja DasGupta

## Abstract

RNA enzymes that assemble RNA from shorter pieces would have been important for primordial RNA-based biology. To explore the scope of RNA-catalyzed RNA assembly, we used directed evolution to re-engineer a ligase ribozyme that uses 5′-phosphorimidazolide RNA substrates into a ribozyme that ligates RNA substrates carrying the biologically relevant 5′-triphosphate group. Unexpectedly, this single RNA evolution experiment generated, in addition to the desired triphosphate ligase, four other distinct classes of ligase ribozymes. Three of these four ligase classes were isolated at extremely low abundances, collectively covering <0.1% of the selected RNA population. One class exhibits strict specificity for 5′-phosphorimidazolide substrates, even though these substrates were never presented during evolution. Another ligase class mediates a reaction between its 5′-triphosphate and the substrate’s terminal 2′-OH, but only when the substrate carries a 3′-phosphate. This reaction resembles those catalyzed by protein enzymes such as RtcB ligase and plant RNA splicing ligases in RNA repair pathways. The remaining classes catalyze two different but related reactions, each involving the 5′-triphosphate groups on the ribozyme and a distinct internal hydroxyl group on the substrate. These reactions resemble branching reactions catalyzed by naturally occurring ribozymes such as the group II intron and the spliceosome, even though these ribozymes emerged from a synthetic RNA library with no relation to biological catalysts. The emergence of such catalytic diversity from a single RNA evolution experiment under highly constrained selection pressures and the discovery of new enzyme reactivities in rare RNA sequences highlight the catalytic flexibility and evolutionary potential of RNA. These findings strengthen the plausibility that an RNA World could have supported a wide range of chemistries essential for the emergence of life.

## INTRODUCTION

Catalytic RNA molecules, referred to as ribozymes, offer crucial insights into the biochemistry of early life before the appearance of proteins. By identifying ribozymes that catalyze reactions relevant to primordial metabolism and genome replication, researchers have begun to reconstruct prebiotically plausible chemical and biochemical pathways to reveal how the first forms of life may have emerged and evolved almost 4 billion years ago.^1–4^ Although ribozyme catalysis is limited to RNA cleavage, RNA ligation, and peptide bond formation in nature, combinatorial methods such as *in vitro* selection and directed evolution have generated a diverse collection of ribozymes that catalyze reactions ranging from nucleotide synthesis, aminoacylation and alkylation of RNA, and templated nucleotide polymerization, to non-biological transformations such as Diels-Alder reactions and aldol condensations.^5–9^ RNA ligation is one of the most extensively studied ribozyme-catalyzed reactions and is especially significant in the context of the RNA World model of the origins of life.^10^ Ligation enables the synthesis of long RNA sequences with the potential to form functional molecules in fewer steps compared to RNA polymerization, and thus provides a more facile pathway to access molecular complexity. Therefore, exploring the catalytic scope of RNA ligase ribozymes is important for reconstructing the RNA-based biochemistry of the RNA World.

RNA ligase ribozymes illustrate the versatility of RNA catalysis. Ligase ribozymes have been evolved that catalyze reactions involving the 2′-hydroxyl, 3′-hydroxyl, or even 5′-phosphate group of one RNA molecule and an activated 5′-phosphate on a second RNA molecule.^7,11,12^ Ribozyme-catalyzed ligation usually involves a reaction between an RNA oligonucleotide substrate and the ribozyme itself, resulting in the covalent modification of the ligase ribozyme. In some cases, these ‘cis-acting’ ribozymes have been converted to ‘trans-acting’ ribozymes by physically separating the catalytic domain of the ligase from the region that directly participates in ligation, although the core reactivities of the ligases remain unchanged.^13,14^ Although several ligase ribozymes have been reported over the past decades, ^15,16^ the limited number of ribozyme selection experiments as well as a limited characterization of their outputs leave open the possibility that some RNA ligase phenotypes remain undiscovered within the vast RNA sequence space. Considering the importance of RNA ligation in the RNA World, it is essential to assess how widely distributed and abundant ligase ribozymes are in sequence space. Also important is determining how many distinct ligation pathways can be supported by RNA, as this will reveal the full extent of RNA’s catalytic prowess.

The catalytic capabilities of RNA may be explored through the directed evolution of existing function, in addition to *de novo* selection from randomized libraries.^17,18^ Previously, we used directed evolution to investigate how ribozymes may switch substrate specificity by evolving a ligase ribozyme that uses RNA substrates bearing the prebiotically relevant 5′-phosphor-2-aminoimidazole (5′-AIP) group to catalyze ligation with substrates 5′-activated with the biologically relevant triphosphate (5′-PPP) group.^19–21^ This was accomplished by challenging a partially randomized library (doped at 21%) composed of AIP-ligase variants with a 5′-triphosphorylated RNA oligonucleotide bait possessing a 3′-TEG-biotin moiety (henceforth, ‘PPPLigB’), followed by capturing the active sequences on streptavidin-coated magnetic beads (Fig. S1). While this experiment yielded the desired ‘triphosphate ligase’ (referred to as CS3) (Fig. 1A) that catalyzes ligation between the ribozyme 3′-OH and the substrate 5′-PPP groups, the other four sequences we characterized (referred to as CS1, CS2, CS4, and CS5) were found to exhibit a novel and unexpected activity (Fig. 1A).^22^ These ribozymes catalyze ligation between the substrate 2′-OH and the ribozyme 5′-PPP only when the substrate possessed a 3′-phosphate (3′-P) group vicinal to the nucleophilic 2′-OH. The sequences comprising the five ribozyme families represented by CS1-CS5 covered >99% of the entire isolated RNA population, indicating their overwhelming fitness for the selection pressures imposed during *in vitro* evolution.

**Figure 1.**
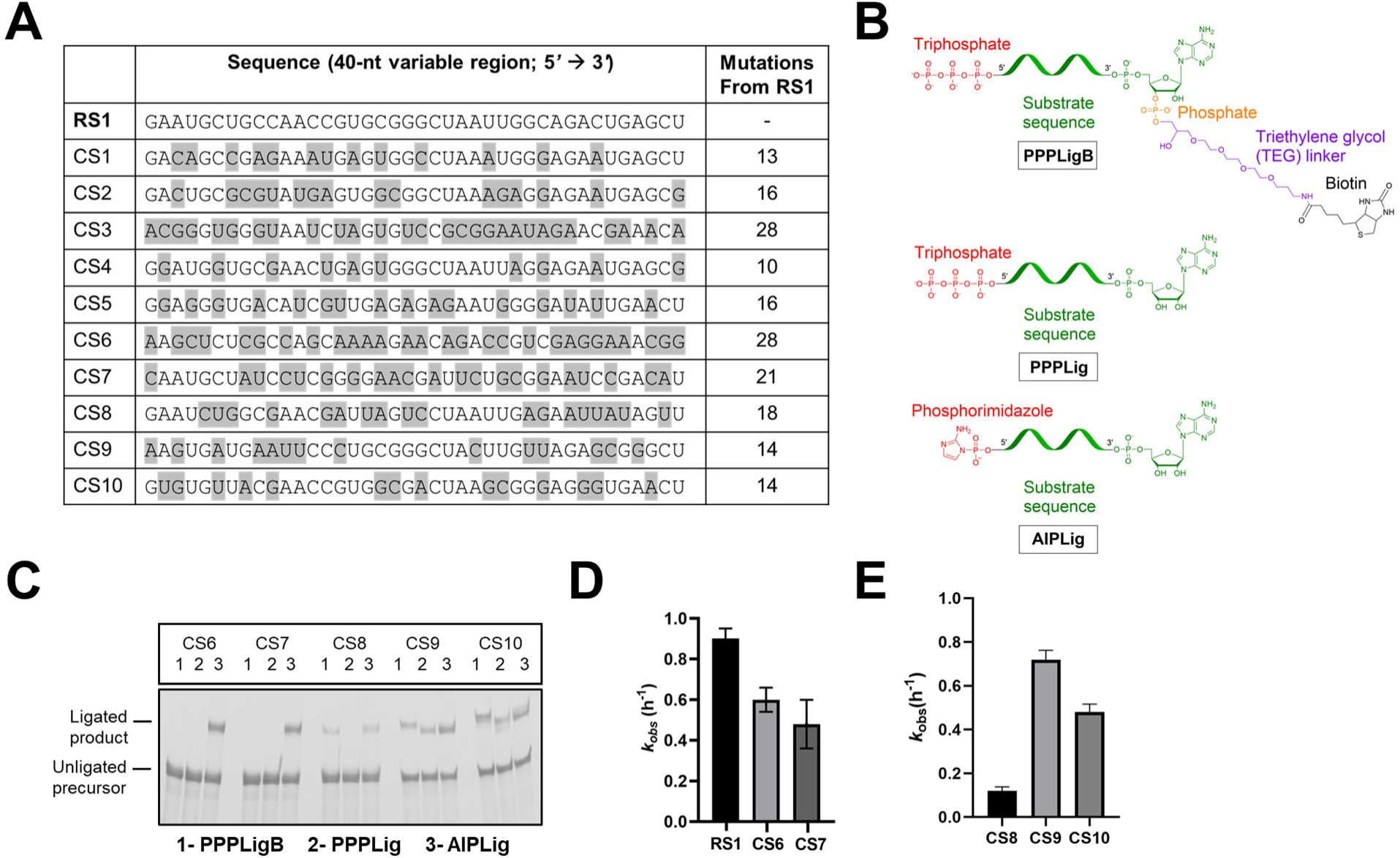
Ligase reactivities of the low-abundance sequences, CS6-CS10. (A) The 40-nt variable region in the dominant sequences of the ten most abundant sequence families, CS1-CS10. RS1 is the ancestral AIP-ligase that CS1-CS10 were evolved from. Sequence divergence from RS1 is highlighted in gray: isolated sequences were 10-28 mutations from RS1. CS1-CS5 have been characterized in earlier publications.^21,22^ Low-abundance sequences, CS6-CS10 were present in 0.01-0.05% abundance in the isolated RNA population. (B) Substrates tested for RNA ligation catalyzed by CS6-CS10. RNA substrates have either triphosphate (PPP) or phosphorimidazole (AIP) groups (red) on their 5′ ends. 5′-PPP substrates have a terminal diol or a 3′-TEG-biotin moiety, which consists of a 3′-phosphate (orange) on the terminal adenine nucleotide, which is connected to a biotin moiety (black) via a triethylene glycol (TEG) linker (purple). (C) CS6-CS10 exhibit diverse reactivities toward RNA substrates containing: 5’ triphosphate and 3’ TEG-biotin groups (PPPLigB), a 5*’* triphosphate group (PPPLig), and a 5′-phosphoro-2-aminoimidazole group (AIPLig). CS6 and CS7 only ligate to AIPLig, CS8 ligates to PPPLigB and, with lesser efficiency, to AIPLig (nonenzymatic ligation), but not to PPPLig. Both CS9 and CS10 ligate to all three substrates. Ligations were assayed at 3 h. (D) CS6 and CS7 exhibit *k*_obs_ values of 0.6 and 0.5 h^−1^ with AIPLig, respectively, making them ∼1.5-fold slower than RS1. (E) CS8, CS9, and CS10 exhibit *k*_obs_ values of 0.1 h^−1^, 0.7 h^−1^, and 0.5 h^−1^, respectively, with PPPLigB, the substrate used in selection. Ligation reactions contained 1 µM ribozyme, 1.2 µM RNA template, and 2 µM RNA substrate (PPPLigB/PPPLig/AIPLig) in 100 mM Tris-HCl (pH 8.0), 300 mM NaCl, and 10 mM MgCl_2_ (for AIPLig kinetics in (D) or 100 mM MgCl_2_ (for PPPLigB/PPPLig/AIPLig ligation assays in (C)). Experiments were performed in triplicate; error bars indicate standard error of the mean (S.E.M).

The isolation of two distinct classes of ligase ribozymes exhibiting different ligase reactivities from a single *in vitro* evolution experiment led us to ask a deeper question: How many different ligase reactivities can emerge from a single evolution experiment? According to practice, we had only focused on the high abundance sequences, but we wondered if sequences isolated at low abundances could harbor unique ligase functions. To explore this possibility, we re-analyzed the high-throughput sequencing data obtained from the directed evolution of the AIP-ligase to PPP-ligases, focusing on sequences in the low-abundance regime. In this work, we report the discovery of five low-abundance ligase sequences, CS6-CS10, representing four different classes of ligase ribozymes that catalyze RNA ligation via distinct chemical pathways. Each of these ribozymes possesses unique properties that add to the catalytic richness of RNA. Two sequences, CS6 and CS7, were found to catalyze ligation exclusively with 5′-AIP-substrates. As the intention of this experiment was to convert an AIP-ligase to a PPP-ligase, no 5′-AIP-substrate was presented during *in vitro* evolution, making the emergence of these new AIP-ligases particularly intriguing. Another ribozyme, CS8, acts as a 3′-phosphate–specific ligase, similar to CS1, CS2, CS4, and CS5, and has implications in primordial RNA repair.^22^ Finally, two sequences, CS9 and CS10, were found to catalyze distinct branching reactions through different internal nucleotides of the substrate, a reactivity reminiscent of the first transesterification step in pre-mRNA splicing – an essential process in extant biology. The isolation of branching ribozymes whose sequences diverge significantly from biological splicing catalysts, and their emergence from a pool with no relation to biological RNAs, suggest that such a vital catalytic reactivity may be supported by independent structural configurations within RNA that are not found in nature. The emergence of five distinct variations of one reaction – ligation – from a single RNA evolution experiment underscores the remarkable evolvability of RNA and its potential to generate biochemical diversity consistent with the hypothetical RNA World.

## RESULTS

### Low-abundance ligase ribozymes isolated from *in vitro* evolution

In our previous work,^21^ we characterized only those sequences whose abundance exceeded an arbitrary threshold of 0.1%, with the expectation that sequences with lower abundances would be significantly less fit and would not show detectable ligase activity. CS1-CS5, the dominant sequences (‘peak sequences’) of the five most abundant ligase families, satisfied this criterion. However, because all but one (CS3) of the isolated sequences exhibited an unexpected, and importantly, novel ligase reactivity, in this work, we investigated the peak sequences of the remaining five ligase families that were detected at a total of ∼0.1% abundance. These sequences, CS6-CS10 (Fig. 1A), each covered only 0.01-0.05% of the isolated sequence population, and diverged significantly from the AIP-ligase (referred to as RS1) they were evolved from (Fig. 1A). This large divergence of 14-28 nucleotides from the ancestral RS1 sequence was unexpected, considering that the library was mutagenized at 21% per position, which generates a pool enriched (by >75%) in variants that are 6-11 mutations from RS1 with only 3% of the starting pool at ≥14 mutations from RS1 (Fig. S2, Table S1). Abundance falls off precipitously with an increase in mutational distance, with 0.04% sequences at 18 mutations and less than 0.0009% sequences at 21 mutations from RS1, respectively. CS6, which occupied 0.05% of the isolated pool, is 28 mutations from RS1, and therefore, was likely only 1 in ∼35,000 sequences in the starting pool of 10^15^ sequences (i.e., 3.5 x 10^-9^ % abundance). The isolation of sequences with extremely low representation in the original selection pool highlights the power of combinatorial selection to selectively amplify rare but fit sequences.

Some sequences can persist through selection-amplification cycles despite weak catalytic activity, owing to their higher amplification efficiency. To verify that CS6–CS10 were, in fact, ligase ribozymes, we tested each sequence for ligase activity using the 5′-triphosphorylated, 3′-biotinylated substrate (PPPLigB) used during *in vitro* evolution (Fig. 1B). Additionally, we tested ligation with a 5′-triphosphorylated substrate (i.e., without the TEG-biotin moiety, but with a terminal diol; henceforth, PPPLig) and a 5′-phosphorimidazolide substrate (henceforth, AIPLig) (Fig. 1B). The inclusion of PPPLig was motivated by our prior discovery that ribozymes, CS1, CS2, CS4, and CS5, catalyze ligation with PPPLigB, but not with PPPLig – a difference that was eventually traced to their dependence on the substrate’s 3′-phosphate group that links its sequence to the TEG-biotin group (Fig. 1B).^22^ Ligation with AIPLig was tested to ensure that the evolved sequences had lost the function of their ancestral sequence, RS1.^19^ Interestingly, we found that CS6-CS10 exhibited different reactivities when challenged with these three substrates (Fig. 1C).

CS6 and CS7 catalyzed ligation with AIPLig but failed to ligate to PPPLigB or PPPLig, despite being evolved to ligate to 5′-triphosphorylated RNA substrates (Fig. 1C). This activity resembles that of AIP-ligase, RS1, with CS6 and CS7 showing ∼1.5-fold slower ligation than RS1 (Fig. 1D, Fig. S3).^23^ CS8 ligated to PPPLigB, but not to PPPLig, thus resembling the 3*’*-phosphate-dependent ligases, CS1, CS2, CS4, and CS5, while showing weak activity for AIPLig, consistent with slow template-directed nonenzymatic ligation at 100 mM Mg^2+^ (Fig. 1C). In contrast, CS9 and CS10 displayed broader substrate reactivity, catalyzing ligation with all three substrates. This insensitivity to the 5′-chemistries of the substrate and to the requirement for a 3′-phosphate group on their substrates is in sharp contrast to the reactivities of CS6-CS8 (Fig. 1C). CS8, CS9, and CS10 showed different ligation efficiencies with PPPLigB, with ligation yields of ∼10, ∼27, and ∼43% after 4 h and *k*_obs_ values of 0.1 h^−1^, 0.7 h^−1^, and 0.5 h^−1^, respectively (Fig. 1E, Fig. S4). In addition to establishing that sequences isolated at extremely low abundance from experimental RNA evolution may harbor catalytic function, the unique reactivities of these rare sequences prompted further investigation to unravel the chemical pathways these ribozymes utilize for catalyzing RNA ligation. In the following sections, we delineate the chemical pathways behind the reactivities of these newly discovered ribozymes, which reveal interesting features about the catalytic diversity of RNA and its evolutionary implications.

### New RNA-based solutions for ligating 5′-phosphorimidazolide RNA

The phosphorimidazole group is considered a prebiotically relevant activating group for the assembly of RNA monomers and oligomers.^19,20^ This makes ribozymes that catalyze ligation with phosphorimidazolide RNA substrates particularly interesting for studying the RNA World. Given that CS6 and CS7 catalyze ligation with substrates containing the 5′-phosphorimidazole group and not the 5′-triphosphate group, similar to the ancestral AIP-ligase, RS1, we considered the possibility that these ribozymes represented conventional template-directed 3′–5′ ligases like RS1 (Fig. 2A).^19^ To investigate this, we tested truncated variants of CS6 and CS7 for ligation with AIPLig, where inactivity of the 3′-truncated construct would support the participation of the ribozyme 3′ end in ligation. We generated 5′-truncated (5′t) constructs by deleting the first 25 nucleotides (shown in orange in Fig. S1) from the 5′ end, and 3′-truncated (3′t) constructs by deleting the last 14 nucleotides from the 3′ end (Fig. 2A). 3′ truncation deletes the hexauridine linker and the 8-nt ‘primer’ (shown in gray and red, respectively in Fig. S1) that is expected to participate in this ligation reaction. 5′-truncated variants of both CS6 and CS7 retained ligase activity, while the 3′-truncated ribozymes lost function completely (Fig. 2B). This suggests that the sequence at their 3′ ends is essential for catalytic activity, either by contributing to the ribozyme’s active conformation or by directly participating in ligation. The inability of CS6 and CS7 to catalyze ligation in the absence of the 16-nt external template, an RNA that bridges the last 8 nucleotides of the ribozyme and the first eight nucleotides of the substrate (Fig. 2C, Fig. S1) supports the direct participation of the ribozyme 3′ end in ligation. To further confirm this reactivity, we next treated the purified products of overnight ligation reactions between each ribozyme and AIPLig with RNase R, a 3′-5′ exonuclease that stalls at noncanonical 2′-5′ phosphodiester linkages (Fig. S5A). Complete digestion of the ligated products generated by CS6 and CS7 (Fig. S5B) suggested the presence of a 3′-5′ phosphodiester bond between the ribozyme and substrate, as RNase R digestion would have produced an RNA that is 1-nt longer than the ribozyme (96-nt) in case of a 2′-5′ linkage. This confirmed that CS6 and CS7 catalyze ligation between their 3′-OH and the substrate’s 5′-AIP groups, a reactivity similar to RS1.^19^ The similarity to RS1 prompted us to investigate whether CS6 and CS7, like RS1, utilize complementary base-pairing with the 3′ end of the substrate, which remains unpaired upon template binding (Fig. S1). Deleting this 8-nt region (5′-AUUCCGCA-3′), referred to as the 3′ overhang, was found to cause a significant decrease in ligation in RS1^19^ and has been shown to be a general catalytic strategy in AIP-ligases.^23^ Both CS6 and CS7 contain regions complementary to the substrate 3′ overhang: in CS6, it is a 4-nt region (5′-GGAA-3′) and in CS7, an 8-nt region (5′-UGCGGAAU-3′). To test the importance of substrate complementarity, we tested ligation by CS6 and CS7 with an 8-nt truncated version of the AIPLig (AIPsLig: 5′-AIP-ACCACCGC-3′) that lacked the 3′ overhang. Mirroring the behavior of RS1, ligation decreased for both CS6 and CS7 with this shorter substrate (Fig. 2D).

**Figure 2.**
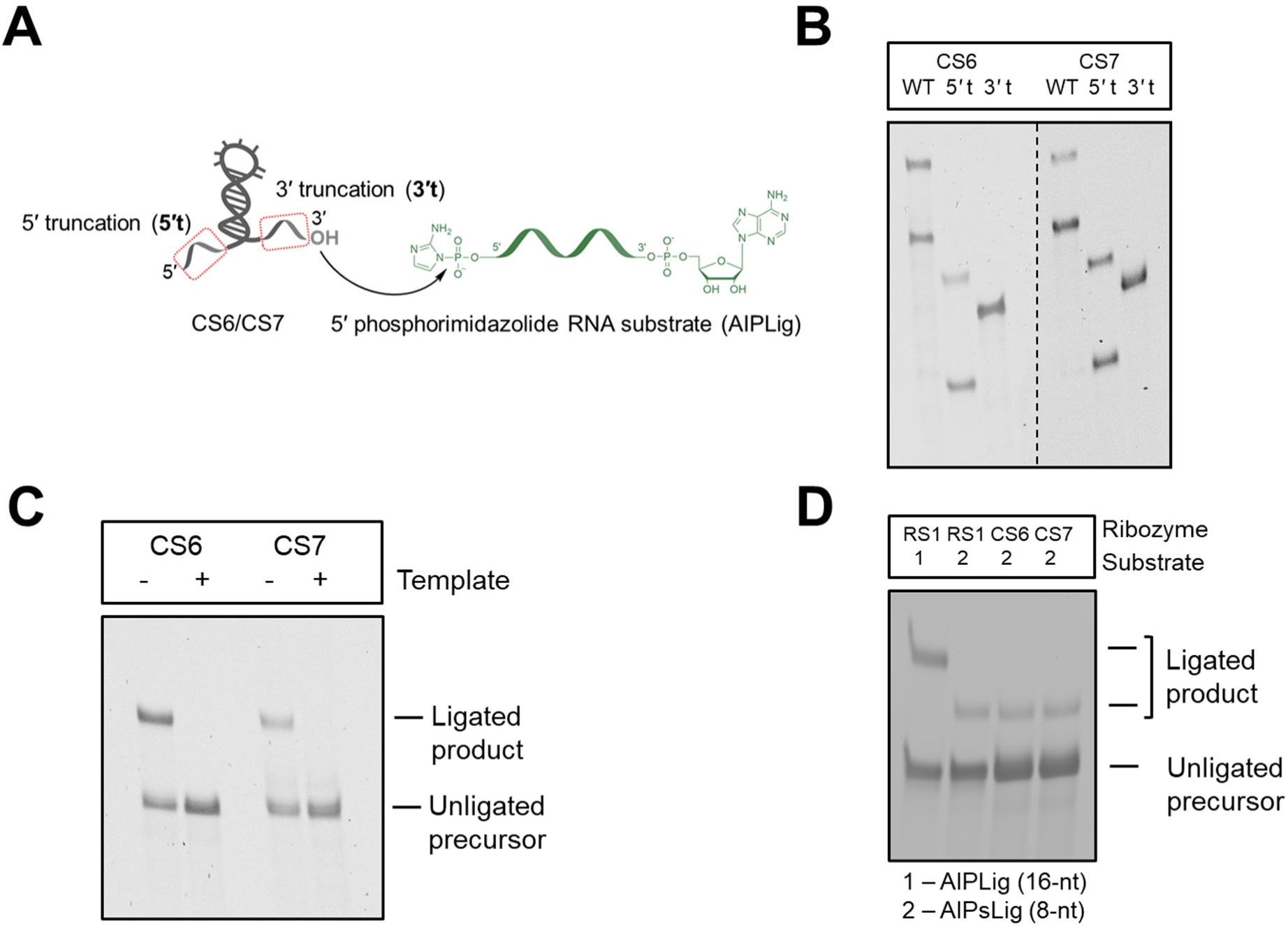
Ligation reaction between the ribozyme 3*’*-end and 5*’*-AIP of the substrate. (A) Schematic for CS6-and CS7-catalyzed ligation; external RNA template not shown. (B) Truncating CS6 and CS7 by deleting the first 25 nucleotides from their 5′ end preserves ligation; however, deleting the last 14 nucleotides from the 3′ end abrogates ligation. (C) CS6-and CS7-catalyzed ligation with AIPLig requires an external template. (D) CS6 and CS7 ribozymes exhibit weak ligation with the eight-nucleotide truncated version of AIPLig (AIPsLig: 5′AIP-ACCACCGC-3′), similar to RS1. Ligation reactions contained 1 µM ribozyme, 1.2 µM RNA template, and 2 µM AIPLig in 100 mM Tris-HCl (pH 8.0), 300 mM NaCl, and 10 mM MgCl_2_. Ligations were assayed at 3 h.

Although CS6 and CS7 resemble RS1 in their reactivities, the three sequences diverge significantly from each other. While CS6 and CS7 are 31 mutations from each other (i.e., they have only 9 conserved nucleotides in the 40-nt variable region), they are 28 and 21 mutations from RS1, respectively (Fig. 1A). Accordingly, their secondary structures determined by SHAPE probing are notably different (Fig. S6). Therefore, RS1, CS6, and CS7 represent distinct solutions to phosphorimidazolide RNA ligation that are distant from each other in sequence space, representing distinct peaks in this fitness landscape.^24,25^

### A new ligase ribozyme that exhibits specificity towards 3′-phosphorylated RNA substrates

The apparent dependence of CS8 on substrate 3′-biotinylation (Fig. 1C) is similar to what was observed for CS1, CS2, CS4, and CS5, which results from the requirement of a 3′-phosphate group that connects the substrate sequence to the TEG-biotin moiety (Fig. 1B).^22^ Like those ligases, CS8 retained activity upon 3′ truncation, albeit at a reduced level, but lost function upon 5′ truncation, suggesting an essential role of the 5′ end (Fig. 3A, 3B). To uncouple any potential structural effects of 5′ truncation from the direct involvement of the ribozyme 5′ end, we tested CS8 variants bearing 5′-phosphate (5′-P) and 5′-hydroxyl (5′-OH) groups, in addition to the ‘wild-type’ 5′-triphosphorylated CS8. CS8 catalyzed ligation only when it carried a 5′-triphosphate, regardless of the substrate’s 5′ end (PPP, P, or OH) (Figs. 3A, 3C, 3D). Ligation with substrates without 5′ activation ruled out the involvement of the substrate 5′ end, and the requirement for a 5′-PPP group on the ribozyme implicated the triphosphate group as the site of nucleophilic attack by the substrate. Most importantly, CS8 catalyzed ligation with substrates bearing 3′-phosphates, but not with those ending in 2′, 3′-diols, a feature similar to CS1, CS2, CS4, and CS5 (Fig. 3A, 3D). Considering the similarities with the previously reported 3′-phosphate-specific ligases, we compared ligation catalyzed by CS8 with CS1, the dominant sequence in this class. CS8 exhibited a rate constant of 0.09 h^−1^ with a 3′-phosphorylated RNA substrate (OHLig3′P), yielding 21% product after 7 hours, making it 10-fold slower than CS1 (Fig. 3E, Figs. S7A and S7B). Like CS1 (and CS2, CS4, and CS5), CS8 failed to catalyze ligation when the substrate’s terminal nucleotide (rA16) was replaced with a deoxyribonucleotide (dA16), confirming the substrate terminal 2′-OH as the nucleophile (Fig. 3F).^22^ The role of the terminal nucleotide of the substrate is further supported by its resistance to mutation in CS8-catalyzed ligations (Fig. S8A). Similar to the others in its class that require a 5′G for function, CS8 with a 5′-terminal G to A mutation is inactive for ligation (Fig. S8B). Collectively, these results revealed the chemical pathway for CS8-catalyzed ligation as a nucleophilic attack of the substrate terminal 2′-OH on the ribozyme 5′-PPP, when the substrate possesses a 3′-phosphate group vicinal to its terminal 2′-OH, generating a 2′-5′ linkage, similar to the reported previous ligases.^22^

**Figure 3.**
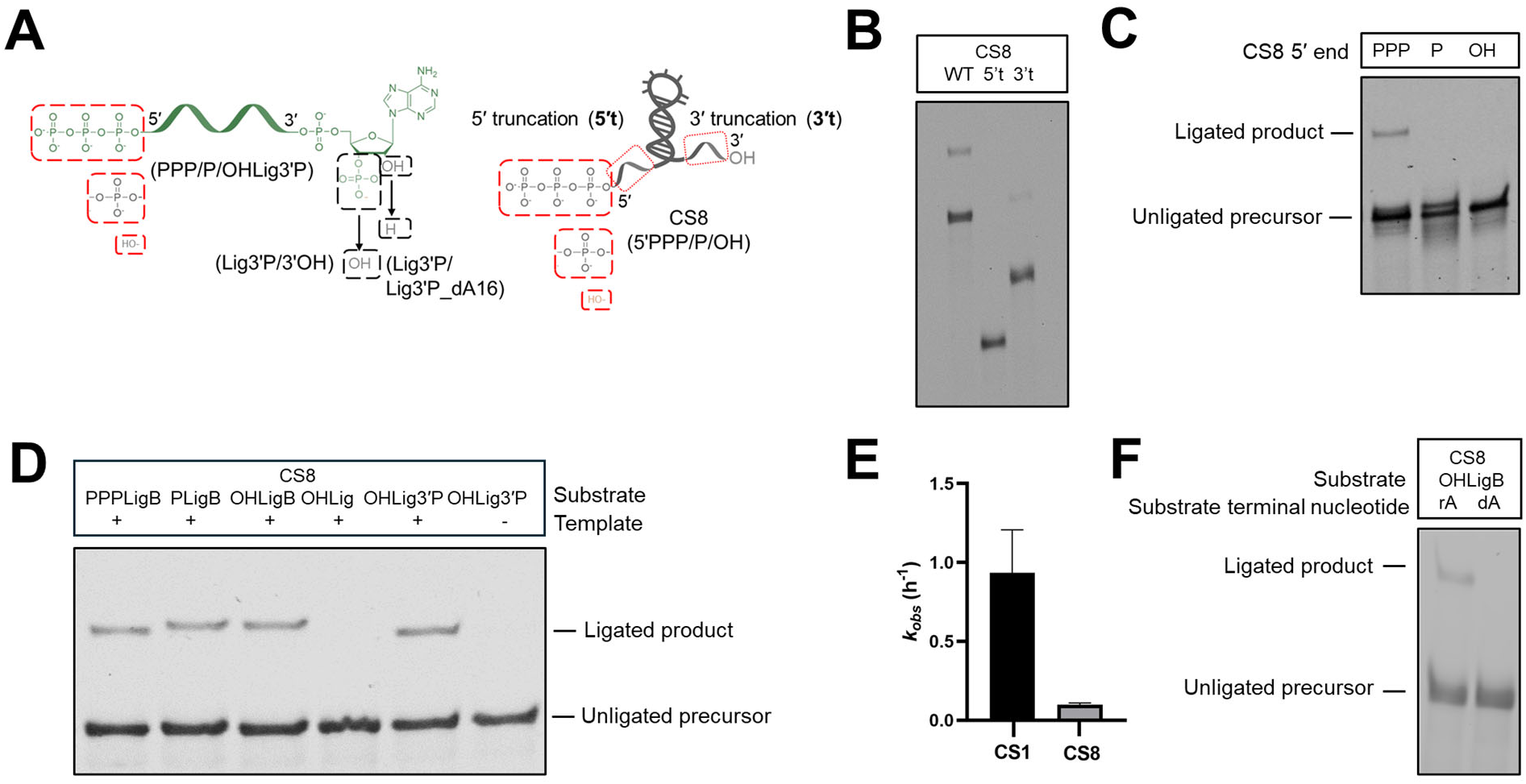
CS8-catalyzed ligation involves the 5′-PPP group of the ribozyme and terminal 2′-OH group of the 3*’*-phosphorylated substrate. (A) Modifications to the substrate and ribozyme to probe the ligation pathway of CS8. (B) Truncating CS8 by deleting the first 25 nucleotides from its 5′ end abrogates ligation; however, deleting the last 14 nucleotides from the 3′ end preserves ligation. (C) Ligation requires a PPP-group at the ribozyme 5′ end. (D) Ligation requires a 3′-phosphate group on the substrate and the presence of an external template. (E) CS8-catalyzed ligation is significantly slower with OHLig3′P compared to CS1, with CS8 and CS1 exhibiting *k*_obs_ values of 0.09 h^−1^ and 0.9 h^−1^, respectively. Experiments were performed in triplicate; error bars indicate standard error of the mean (S.E.M). (F) Ligation by CS8 was abolished when the 3′ terminal adenine of the substrate was changed to deoxyadenine, indicating the importance of its 2′-OH group in ligation. Ligation reactions contained 1 µM ribozyme, 1.2 µM RNA template, and 2 µM substrate in 100 mM Tris-HCl (pH 8.0), 300 mM NaCl, and 100 mM MgCl_2_.

In our previous study, we found that of the 3′-phosphate-specific ligases, only CS1 was active without an external template. This template dependence stems from base-pairing between the ribozyme’s 3′ end sequence (‘primer’) and the template (Fig. S1 and S9A). CS8 exhibited a similar template dependence (Fig. 3D). Disrupting this interaction by mutating the template sequence abolished ligation, which was restored by compensatory mutations to the primer sequence (Figs. S9A, B). This result is consistent with the reduced activity of the 3′-truncated version of CS8, which lacks the 3′ primer (Fig. 3B). By forming base-pairs with the ribozyme 3′ end and substrate 5′ end, the template potentially helps to position the substrate close to the active site of the enzyme. SHAPE probing of CS8 suggests a secondary structure where the 3′ primer remains unpaired and distal from the rest of the ribozyme separated by the U_6_ linker, allowing base-pairing interactions with an external template (Fig. S10). As expected from the sequence divergence of 18 nucleotides from CS1, the two ligases adopt distinct secondary structures.

### Ribozymes that catalyze RNA branching

The ability of CS9 and CS10 to catalyze ligation with substrates containing phosphorimidazole or triphosphate groups at their 5′ ends (Fig. 1B, C) indicated that either the 5′ end of the substrate was not involved in ligation or these ligases were catalytically promiscuous. The possibility of promiscuity was eliminated when we found that unactivated RNAs with 5′-P or 5′-OH groups were also substrates for ligation (Fig. 4A, B). The loss of function in 5′-truncated versions (5′t) of CS9 and CS10 when incubated with PPPLigB supported the involvement of their 5′ ends (Fig. 4A, C). Deleting the last 14 nucleotides from the ribozyme 3*’* ends (3′t) eliminated ligation for CS9, but preserved ligation for CS10 (Fig. 4A, C). The loss of function in the 3′-truncated version of CS9 was found to be due to the loss of base-pairing interactions between the 3′ end sequence of CS9 (‘primer’) and the external template (Fig. S9A). Ligation defects due to the disruption of this interaction through template mutations were rescued through compensatory mutations in the ribozyme 3′ primer sequence (Figs. S9A, C). This suggests a beneficial role of the template in positioning the substrate suitably for catalysis by base pairing with the ribozyme 3′ end and substrate 5′ end. This finding aligns with this ribozyme’s reliance on an external template and CS10’s ability to ligate without one (Fig. 4B). A complete loss of activity in CS9 and CS10 variants containing 5′-P or 5′-OH groups bolstered the involvement of the ribozyme 5′-PPP group in ligation (Fig. 4A, D).

**Figure 4.**
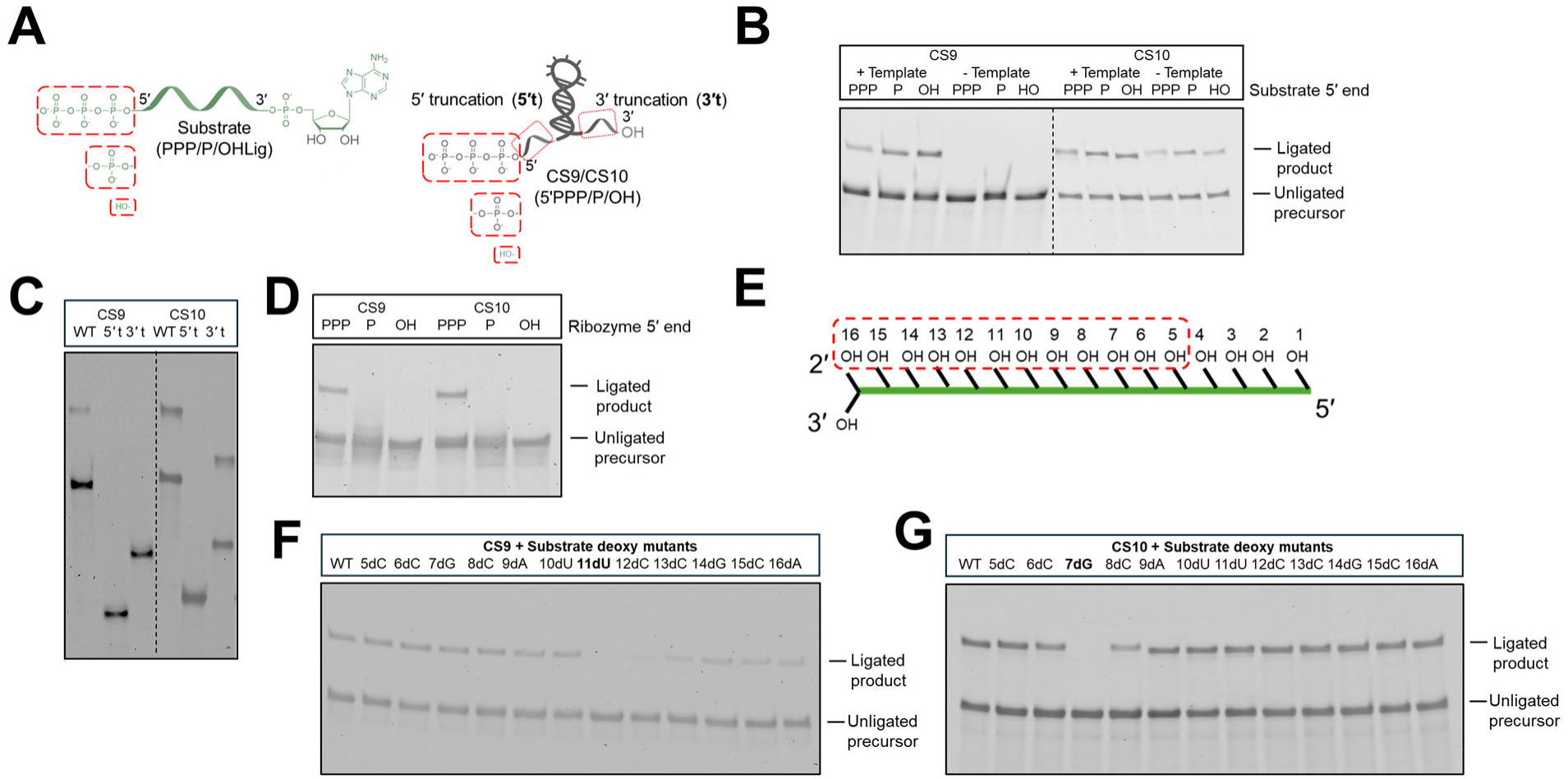
Ligation proceeds through a branching reaction that involves an internal 2*’*-OH of the substrate and the 5′ PPP of the ribozyme. (A) Modifications to the substrate and ribozyme to probe the ligation pathways of CS9 and CS10. (B) Ligation catalyzed by CS9 and CS10 is agnostic about substrate 5′ chemistry. CS9 needs a template for ligation; however, ligation by CS10 is template-independent. (C) Truncating CS9 and CS10 by deleting the first 25 nucleotides from their 5′ ends abrogated ligation for both CS9 and CS10. Deleting the last 14 nucleotides from the 3′ end eliminated ligation for CS9, but preserved ligation in CS10. (D) Ligation proceeds through the 5′ PPP groups of CS9 and CS10. (E) Internal 2*’*-OH groups in the substrate RNA that were subject to deoxy substitution. (F) A substrate containing a deoxyribonucleotide at the 11^th^ position fails to ligate to CS9, establishing the internal 2*’*-OH of U11 as the nucleophile for CS9-catalyzed RNA ligation. (G) A substrate containing a deoxyribonucleotide at the 7^th^ position fails to ligate to CS10, establishing the internal 2*’*-OH of G7 as the nucleophile for CS10-catalyzed ligation. Ligation reactions contained 1 µM ribozyme, 1.2 µM RNA template, and 2 µM substrate in 100 mM Tris-HCl (pH 8.0), 300 mM NaCl, and 100 mM MgCl_2_. Ligations were assayed at 3 h.

With the 5′-PPP group of CS9 and CS10 identified as the electrophilic center, and the 5′ chemistry of the substrate established as unimportant for ligation, we anticipated that the 2′-OH group of the 3′ terminal nucleotide of the substrate would serve as the nucleophile in CS9-and CS10-catalyzed ligation – in line with previously isolated ribozymes CS1, CS2, CS4, and CS5.^22^ However, unlike those ligases, CS9 and CS10 ligated substrates modified with a terminal deoxyadenine (dA16) nucleotide (Fig. S11A) indicating that the nucleophile was located elsewhere on the substrate. To hunt for the nucleophile, we substituted all ribonucleotides from the 5^th^ to the 15^th^ positions of the substrate with deoxynucleotides, one at a time (Fig. 4E). Deoxy-substitutions at positions 11 (uridine) and 7 (guanine) on the substrate eliminated ligation for CS9 and CS10, respectively (Fig. 4F, G). This suggests that CS9 activates U11 and CS10 activates G7 – two different nucleotides in the same substrate – to catalyze branching reactions through their internal 2′-OH groups acting as nucleophiles. The nucleotides at the ligation junction – the 5′ terminal G of the ribozymes and the internal U11 or G7 of the substrate – are conserved. CS9 and CS10 largely lost their activities when their 5′ terminal guanines were mutated to adenines (Fig. S11B), with CS10 and CS10_5′A showing weak ligation with the G7A substrate mutant (HOLig_A7) and the wild-type substrate (HOLig), respectively. Substrates with U11 mutated to A, C, or G were not ligated by CS9, and substrates with G7 mutated to C or U were not ligated by CS10 (Fig. S11B). While CS10 was not influenced by any other deoxy mutations to the substrate, a dU12 mutation to the substrate reduced ligation by CS9 (Fig. 4F). A faint ligation band for this mutant suggests that although U12 is not the nucleophile, it plays a role by potentially stabilizing the unusually strained ligation junction required for RNA branching or even activating the U11 nucleophile. The branching reactions catalyzed by CS9 and CS10 resemble the first step of pre-mRNA splicing, wherein a branched adenosine attacks the 5′ splice site, resulting in the loss of the upstream exon and the formation of an intron lariat (Fig. 5D-F).^26^ CS9 and CS10 represent the first ribozymes to catalyze branching derived from a completely non-biological context, and therefore, warrant more detailed biochemical characterization, which will be reported separately.

**Figure 5.**
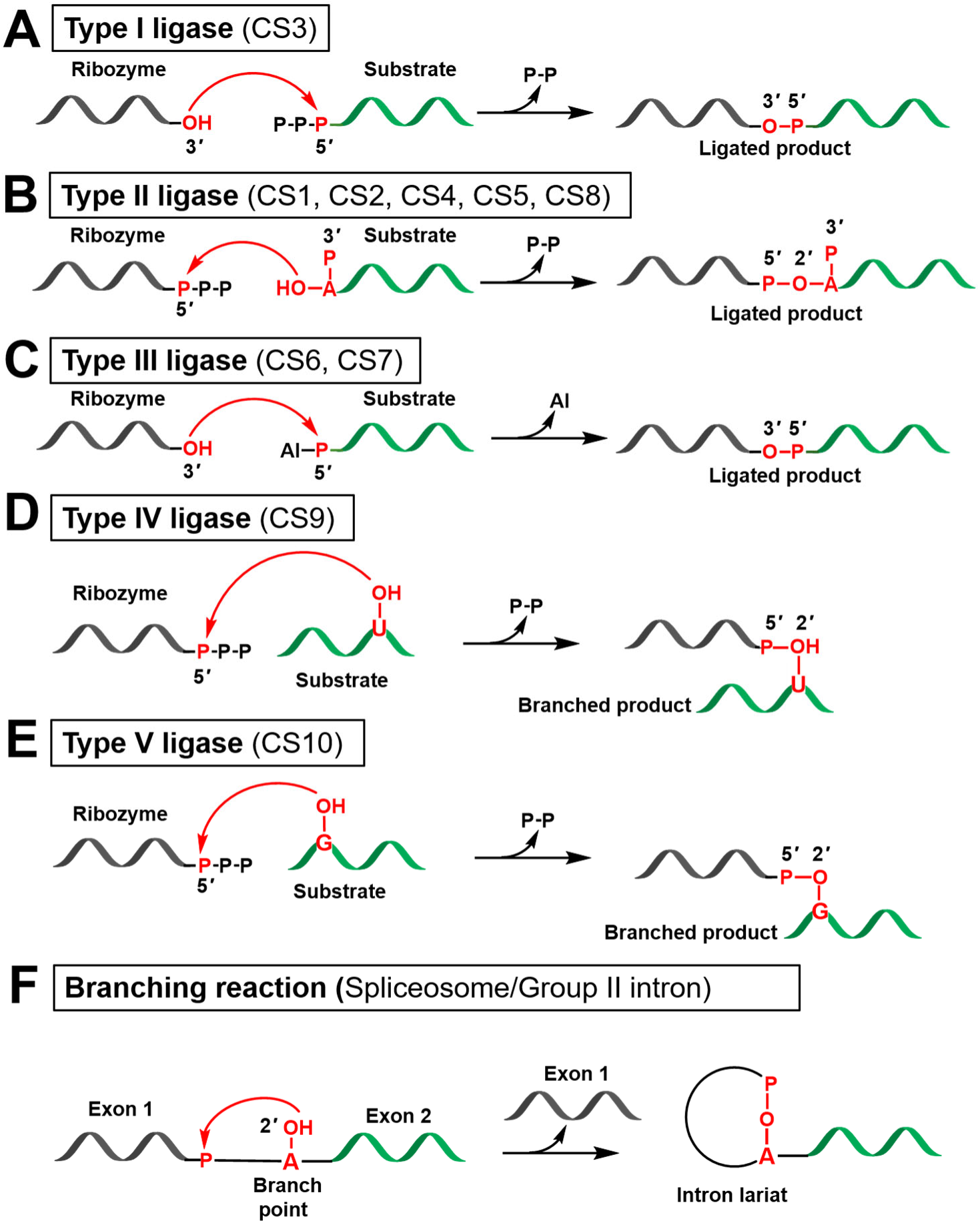
Diverse ligase reactivities that emerged from a single RNA evolution experiment (A-E) and biological branching. (A) Type I ligases catalyze a reaction between the ribozyme 3′-OH and the 5′-PPP of the substrate. This ribozyme also ligates with AIPLig.^21^ (B) Type II ligases catalyze a reaction between the 2′-OH of the substrate and the ribozyme 5′-PPP, in the presence of 3′-phosphate in the substrate.^22^ (C) Type III ligases catalyze a reaction between the 3′-OH of the ribozyme and the 5′-AIP of the substrate (this work). (D) Type IV ligases catalyze a reaction between the 2′-OH of the substrate U11 and the 5′-PPP of the ribozyme creating a branched product (this work). (E) Type V ligases catalyze a reaction between the 2′-OH of the substrate G7 and the 5′-PPP of the ribozyme creating a branched product (this work). (F) Branching reactions similar to Type IV and Type V ligases are catalyzed by naturally-occurring ribozymes that result in the formation of lariat RNA.

### Emergence of five distinct RNA ligase reactivities in a single evolution experiment

A partially randomized RNA library obtained by mutagenizing the 40-nt catalytic core of an *in vitro* selected AIP-ligase, RS1, yielded five distinct ligase reactivities from an experiment designed to isolate PPP-ligases. While Tuschl and Bartel demonstrated more than 25 years ago that multiple ribozymes can emerge from a single *in vitro* evolution experiment,^27^ this work is, to our knowledge, the only example where multiple ribozymes that catalyze a single type of reaction (here, RNA ligation) via distinct catalytic pathways emerged from a single experiment. We isolated a total of ten ribozyme classes from this evolution experiment whose peak sequences, CS1 – CS10, are 10-28 mutations from RS1. It is interesting that, while only ∼32% of sequences in the starting library were 10 or more mutations from RS1, ∼100% of the isolated sequences were ≥10 mutations from RS1 (Fig. S2, Table S1). This increase in mutational distance from the ancestral sequence during directed evolution presumably stems from the need to escape the ancestral structural fold in order to encounter new functions.^28^ This is consistent with our SHAPE data, which revealed that the ligases evolved from RS1 have distinct secondary structures that diverge from the structure of RS1 (Figs. S6, S10).^19,21,22^

CS1, the most abundant sequence isolated at ∼24% abundance, is 13 mutations from RS1, when the total abundance of sequences at this mutational distance in the starting library was just ∼3%. In contrast, CS4, a ligase with the same reactivity as CS1 and 10 mutations from RS1, was present at only 0.2% in the final selected population when 12% of the starting population were 10 mutations from RS1. In some cases, ribozymes with identical or related reactivities were found at the same mutational distance from RS1, although their sequences diverge from each other. For example, CS9 and CS10 – both branching ligases – are at a mutational distance of 14, and CS2 and CS5 – both 3′-phosphate-dependent ligases – are at 16 mutations from RS1. Curiously, CS6 and CS7, ligases with the same reactivity as RS1, are 28 and 21 mutations from it. Their selection indicates that one or a few copies of such rare sequences in the starting library were able to survive the initial rounds of selection-amplification before getting enriched.

The current study, in combination with our prior reports on this *in vitro* evolution experiment, allows us to classify the 10 isolated ligase ribozymes according to their reactivities (Fig. 5 and Table S2). We previously reported CS3 as the desired PPP-ligase, which also exhibits AIP-ligase activity, making it catalytically promiscuous.^21^ This ligase catalyzes a reaction between the ribozyme 3′-OH and the substrate 5′-PPP (or 5′-AIP) in the presence of an external template that bridges the reacting parts of both RNAs (Fig. 5A). We call this family of ribozymes, ‘Type I’ ligases. CS1, CS2, CS4, CS5, and CS8 all catalyze ligation between the substrate 2′-OH and the ribozyme 5′-PPP, but only when the substrate carries a 3′-phosphate (Fig. 5B). Despite their shared reactivity, they differed in template requirement and were thus sub-classified into ‘Type IIa’ (template-independent: CS1) and ‘Type IIb’ (template-dependent: CS2, CS4, CS5, CS8). CS6 and CS7 are AIP-ligases, selected unexpectedly from an evolution campaign designed to diverge from the AIP-ligase phenotype. These ligases catalyze a reaction between the ribozyme 3′-OH and the substrate 5′-AIP, generating a 3′-5′ bond, a reactivity similar to the ancestral ligase, RS1. These ribozymes were classified as ‘Type III’ ligases (Fig. 5C). CS9 and CS10 catalyzed distinct branching reactions that involve the nucleophilic attack of internal 2′-OH groups of the substrate on the 5′-PPP group of the ribozyme. Although CS9 and CS10 use identical substrates, they catalyze branching from different nucleotides in it: U11 for CS9 and G7 for CS10, making their ligation pathways related but distinct (Figs. 5D, E). Additionally, CS9 requires an external template for ligation (Fig. 4B), while CS10 does not. Therefore, we categorized the CS9-family ribozymes as ‘Type IV’ and the CS10-family ribozymes as ‘Type V’ ligases.

## DISCUSSION

The preponderance of catalytic function in the RNA sequence space lies at the foundation of the RNA World model of the origins of life. RNA-based life would not survive if ribozymes were rare and essential reactions such as RNA assembly were chemically constrained. Our initial goal was to evolve new substrate specificity in RNA ligase ribozymes by isolating new triphosphate ligases from a pre-existing phosphorimidazolide ligase. In a curious turn of events, this directed evolution experiment yielded not one, but five different types of ligase that catalyze RNA ligation via five completely distinct pathways, that collectively involve the ribozyme 3′ and 5′ ends, the substrate 2′ and 5′ ends, and even two distinct internal nucleotides on the substrate, distinguishing this outcome from previous instances where related ligases that catalyze 2′-5′ and 3′-5′ templated RNA ligation were isolated.^11,29^

In addition to the target triphosphate ligase (Type I ligase: CS3), we isolated ligases with a unique reactivity: specific ligation to substrates bearing 3′-phosphate groups (Type II ligase: CS1, CS2, CS4, CS5, and CS8). Here we demonstrate that CS8, although isolated at low abundance, exhibits the same reactivity as the more abundant CS1, CS2, CS4, and CS5 despite it being 20-26 mutations away from these ligase sequences. As suggested by SHAPE probing, the catalytic fold of CS8 likely diverges from these ligases; therefore, CS8 represents another unique solution to 3′-phosphate-specific RNA ligation. The ability of various RNA configurations to support this catalytic reactivity is significant in the light of its implication in primordial RNA repair in an RNA World.^22^

This experiment also yielded new phosphorimidazolide ligases (Type III: CS6 and CS7) with distinct sequences and structures, unrelated to the ancestral ligase, RS1, although no phosphorimidazolide RNA substrate was supplied in the selection. This raises the obvious question: how did these ligases get selected if they do not catalyze ligation with 5′-triphosphorylated RNA substrates? While we do not have a clear answer, we can speculate that these sequences have a low level of PPP-ligase activity that remained undetected in our experiments, but was stimulated in the presence of other sequences in the library during evolution. There is precedent for this in the work of Attwater and Holliger, where the authors observed that the weak activities of certain dominant sequences isolated from *in vitro* evolution were stimulated when a different sequence isolated from the same experiment was added to them.^30^ This functional rescue was determined to be due to stabilizing interactions between the two RNA molecules. Such interactions may have been mutualistic, resulting in increased fitness for both sequences involved. Consequently, CS6 and CS7 do not exhibit the ability to catalyze triphosphate ligation independently.

The branching ligases that emerged at the lowest abundance (0.01-0.02%) are intriguing because their reactivity resembles that of naturally-occurring ribozymes, despite being derived from a purely synthetic RNA population. The best example is in pre-mRNA splicing, which is catalyzed by the spliceosomal U2/U6 snRNA complex in eukaryotes and the group II intron ribozyme in prokaryotes (Fig. 5F).^31,32^ In the spliceosome, the U2 snRNA pairs with the intron branch site to position the branch-point adenosine for nucleophilic attack on the upstream exon to generate a 2*’*-5*’*-linked lariat, while the U6 snRNA provides the catalytic core and binds catalytic Mg^2+^.^33^ The group II self-splicing intron also catalyzes the nucleophilic attack of the intron branch site adenosine, leading to the formation of an intron lariat, and is known to release pyrophosphate instead of a 5*’* exon in certain instances.^34,35^ A group-I-intron-like ‘lariat-capping ribozyme’ catalyzes a branching reaction between residues separated by only a single nucleotide, which results in the formation of a 3-nt lariat and the release of the remaining ribozyme sequence.^36^ The 3*’*-truncated form of CS10, standing at 81-nt with potential for further minimization, represents a simpler RNA-only motif with branching activity, relative to the group II intron (400-600 nt) and the lariat-capping ribozyme (∼200 nt).

Tuschl and Bartel evolved branching ribozymes that facilitate ligation between an oligonucleotide substrate and the ribozyme, with the release of a 5*’*-pyrophosphate from a >200-nt library derived (doped at only 3% per position) from the core sequences of the U2/U6 snRNA complex.^27^The proximity of the library sequences to the catalytic RNA sequences of the spliceosome heavily biased this experiment toward the isolation of sequences with the branching phenotype, so much so that the selected branching ribozymes even possessed a conserved ACAGAGA region, a highly conserved sequence in U6 snRNA, which plays a critical role in splicing.^37^ It is interesting to note that although CS9 and CS10 catalyze RNA branching like these ribozymes, they emerged from a library with no connection to biological RNAs, suggesting that catalytic RNA reactivities essential for life can emerge independent of biological contingency. However, both CS9 and CS10 possess a ‘ACA*<u>C</u>*AGA’ sequence in the 5*’* constant region, which is only a single mutation away from the conserved ‘ACA*<u>G</u>*AGA’ in biological RNAs with branching activities. Although this sequence resides within the 5*’* constant regions of the ribozymes and therefore, did not emerge as a result of selection pressure, these branching ligases may have evolved to utilize this sequence in their catalytic strategy. Regardless of the catalytic mechanism, RNA branching appears to be supported by diverse RNA sequences.

One can speculate that ribozymes encoded in viral genomes with similar reactivity to CS9 and CS10 would be useful for capping the immunogenic 5*’*-triphosphate ends of viral RNAs to evade the host immune system. Such a mechanism would serve a similar purpose as ‘cap-snatching’ from host mRNAs^38^ or noncanonical capping with the flavin adenine nucleotide (FAD) moiety.^39^ On the other hand, RNA capping with better leaving groups than pyrophosphate, such as m^7^G, and noncanonical caps like NAD, FAD, and dephospho-CoA^40^ may make cellular mRNAs susceptible to unwanted reactions if they encode branching ribozymes. This would require evolutionary pressures to prevent the selection of RNA motifs that catalyze intramolecular branching. The branching reactivity is not limited to RNA. Work by the Silverman lab has identified DNAzymes that catalyze RNA branching, with some of these enzymes mediating lariat formation with a release of an oligonucleotide leaving group.^41–45^ Our branching ligases add to the collection of nucleic acid-based branching catalysts.

The discovery of four distinct ligase ribozyme reactivities at a total abundance of ∼0.1% highlights the importance of characterizing sequences that emerge with low read counts from combinatorial selections. Before the age of high-throughput sequencing, rare sequences such as these would remain undetected. From the time practitioners of combinatorial selections started using high-throughput sequencing to identify isolated sequences, the general approach has been to cluster related sequences into families and experimentally characterize the representative ‘peak’ sequences of the most abundant families.^19,21,46^ Although this approach is practical as it samples the fittest (and presumably, the most active) sequences while conserving time and resources, our work shows that it may miss low-abundance sequences that harbor unexpected, and even new reactivities. The emergence of five variants of RNA-catalyzed RNA ligation from a single experiment also challenges the notion that ribozymes are rare in the sequence space.^21,47,48^ Ribozymes that catalyze RNA ligation – a vital reaction in extinct RNA-based biology – via multiple distinct pathways, including some that resemble important reactions in extant biology, underscore the versatility of RNA catalysis and strengthen the prospects of an RNA World.

## MATERIALS AND METHODS

### Materials

All the chemical reagents were sourced from Sigma-Aldrich, unless stated otherwise. Enzymes and molecular biology reagents were primarily obtained from New England Biolabs. Phenol-chloroform-isoamyl alcohol and SYBR Gold Nucleic Acid Gel Stain were purchased from ThermoFisher Scientific, while 100% ethanol was acquired from Decon Laboratories, Inc. QIAquick PCR purification kits were sourced from Qiagen. Denaturing polyacrylamide gels were prepared using Sequagel-UreaGel concentrate and its corresponding diluent system, both obtained from National Diagnostics. A complete list of oligonucleotides used in this study is provided in Table S3. Except for PPPLig and PPPLigB, which were procured from Chemgenes, all oligonucleotides were sourced from Integrated DNA Technologies (IDT) or generated by *in vitro* transcription of corresponding DNA templates.

### RNA preparation and substrate activation

Double-stranded DNA (dsDNA) templates generated by PCR, incorporating 2′-O-methyl modifications at the final two nucleotides of the template strand, were used for in vitro transcription to obtain the ribozyme constructs. Each 1 mL transcription reaction contained 40 mM Tris-HCl (pH 8.0), 2 mM spermidine, 10 mM NaCl, 25 mM MgCl_2_, 10 mM dithiothreitol (DTT), 30 U/mL murine RNase inhibitor, 2.5 U/mL thermostable inorganic pyrophosphatase (TIPPase), 4 mM of each NTP, 30 pmol/mL DNA template, and 1 U/μL T7 RNA polymerase. Reactions were incubated at 37 °C for 3 h. Residual DNA template was removed by treatment with 5 U/mL DNase I by incubating at 37 °C for 30 min, followed by RNA extraction using phenol-chloroform-isoamyl alcohol. The RNA was subsequently precipitated with ethanol and further purified by 10% denaturing polyacrylamide gel electrophoresis (PAGE).

CS8, CS9, and CS10 variants featuring modified 5′ termini (5′-P or 5′-OH) were generated via splinted ligation of three RNA oligonucleotide fragments (Table S3). The first fragment carried the 5′ modification: 5′-triphosphate (5′-PPP), 5′-monophosphate (5′-P), or 5′-hydroxyl (5′-OH), while the second and third fragments were 5′-monophosphorylated to facilitate ligation by T4 RNA ligase 2. 1.2 nmol of each RNA fragment was mixed with 0.8 nmol of each of two complementary DNA splints, heated at 90 °C for 3 min, then incubated at 30 °C for 10 min. 1 U/μL of RNA ligase 2 and 1X T4 RNA ligase buffer were subsequently added, and the 20 μL reaction was incubated at 30 °C for 2 h. Following ligation, RNA products were purified using the Zymo Oligo Clean & Concentrator kit per the manufacturer’s protocol, and further resolved by 10% denaturing PAGE.

5′-monophosphorylated oligonucleotides corresponding to the AIP-substrates (Table S3) were activated as previously described. Briefly, it was reacted with 0.2 M 1-ethyl-3-(3-dimethylaminopropyl) carbodiimide (HCl salt) and 0.6 M 2-aminoimidazole (HCl salt, pH adjusted to 6) in aqueous solution for 3 h at room temperature. Salts were removed by four or five washes (200 µL water per wash) in 3kDa cut-off Amicon Ultra spin columns. This was followed by reverse-phase analytical HPLC purification using a gradient of 98% to 89% 20 mM TEAB (triethylamine bicarbonate, pH 8) versus acetonitrile over 40 min.

### Ribozyme-catalyzed RNA ligation assays

Ligation reactions were performed at room temperature using 1 µM ribozyme, 1.2 µM RNA template, and 2 µM RNA substrate in a buffer containing 100 mM Tris-HCl (pH 8.0), 300 mM NaCl, and, unless otherwise noted, 100 mM MgCl_2_. At designated time points, reaction aliquots were quenched with 5 volumes of quench buffer (8 M urea, 100 mM Tris-Cl, 100 mM boric acid, 100 mM EDTA) and analyzed by 10% denaturing PAGE. Gels were stained with SYBR Gold and imaged using the Amersham Typhoon RGB scanner (Cytiva). Image analysis was carried out in ImageQuant IQTL 8.1. Kinetic data were fitted to a modified first-order rate equation, y = A (1 – e^−*k*x^), where A is the fraction of active complex, *k* is the first-order rate constant, x is time, and y is the fraction of ligated product, using GraphPad Prism 9.

### Confirmation of 3′-5′ phosphodiester bonds in the ligated products generated by CS6 and CS7

∼4 pmol of the purified ligation product, generated from an overnight reaction between CS6/CS7 and AIPLig, was treated with 20 U of RNase R exonuclease (Lucigen) in a 10 μL reaction containing 1X RNase R buffer. The digestion was carried out at 50 °C for 2 h. The reaction was quenched by adding 0.3 μL of 0.5 M EDTA, followed by heat inactivation of the enzyme at 95 °C for 3 min. Subsequently, 10 μL of quench buffer (8 M urea, 100 mM Tris-Cl, 100 mM boric acid, 100 mM EDTA) was added, and the products were resolved by 10% denaturing PAGE.

### Secondary structure determination by SHAPE probing

SHAPE probing of CS6, CS7, and CS8 was performed following the protocol described by Walton *et al.*, 2020.^19^ Briefly, 100 pmol of each RNA construct containing 5′ and 3′ SHAPE cassettes (CS6_SHAPE, CS7_SHAPE, and CS8_SHAPE; Table S3) was folded in 100 mM Tris-HCl (pH 8.0), 250 mM NaCl, and 10 mM MgCl₂, then divided into ‘modification’ and ‘control’ reactions. Approximately 40 mM of the SHAPE reagent 1M7 (dissolved in DMSO) was added to the modification reaction, while an equal volume of DMSO was added to the control. Modified and unmodified RNAs were reverse transcribed using 40 pmol of a 5′ FAM-labeled primer (SHAPE_RT_primer; Table S3) and Superscript III reverse transcriptase (Invitrogen). Sequencing ladders were generated by reverse transcription of 30 pmol unmodified RNA in the presence of each of the four ddNTPs and 25 pmol SHAPE_RT_primer. Quenched reactions were analyzed by 10% denaturing PAGE. Normalized SHAPE reactivities were calculated by excluding the most reactive nucleotide and dividing each reactivity value by the average of the top 10% most reactive positions. These normalized reactivities were then used as constraints for secondary structure prediction in the RNAstructure program.^49^

## ACKNOWLEDGEMENTS

This material is based upon work supported by the National Science Foundation (NSF) under Award No. MCB-2540950. This work was also supported by the University of Notre Dame start-up funds to S.D. and the University of Notre Dame Bioengineering and Life Sciences Postdoc Professional Development Award to A.B. We thank Zoe Weiss for help with bioinformatics analysis of the high-throughput sequencing data in this work. We thank Professor Dipankar Sen, and members of the DasGupta lab – Natalie Kotlin, Ziyao Zhu, Taylor Opolka, Nicholas Colorito, and Dr. Priyanka for helpful comments on the manuscript. We thank Ziyao Zhu for pointing out a critical error that would have otherwise led to erroneous interpretations. We thank Professors Jack W. Szostak and Joseph A. Piccirilli for useful advice early on in the project.

## CONTRIBUTIONS

A.B. and S.D. designed research; A.B. performed research; A.B. and S.D. analyzed the data and wrote the paper.

## DATA AVAILABILITY

The main data supporting the findings of this study are available within the article and additional supporting data for this article have been included in the Supplementary Information file. Raw sequencing data obtained from the in vitro evolution relevant to this work can be accessed at CurateND (https://doi.org/10.7274/30810947). Additional details on datasets and protocols that support the findings of this study will be made available by the corresponding author upon reasonable request.

## CODE AVAILABILITY

The codes used for pre-processing and analyzing sequencing data can be freely accessed on the lab’s GitHub (https://github.com/dasguptalab/Directed-evolution-from-PI-to-PPP-Ligases).

