## Supplementary Information for "Emergence of diverse ligase reactivities from a single RNA evolution experiment"

**This PDF file includes:**

Supplementary Tables 1 to 3

Supplementary Figures 1 to 11

References 1-3

**Table S1. Representation of sequence variants of the ancestral AIP-ligase, RS1, in the starting RNA library mutagenized at 21%.**

| Mutations | Probability (%) | Mutations | Probability (%) |
|---|---|---|---|
| 0 (RS1) | 0.008036809 | 21 | 0.000870415 |
| 1 | 0.085454679 | 22 | 0.000199825 |
| 2 | 0.442958114 | 23 | $4.16 \times 10^{-5}$ |
| 3 | 1.491479219 | 24 | $7.83 \times 10^{-6}$ |
| 4 | 3.667339724 | 25 | $1.33 \times 10^{-6}$ |
| 5 | 7.0190097 | 26 | $2.04 \times 10^{-7}$ |
| 6 | 10.88390745 | 27 | $2.81 \times 10^{-8}$ |
| 7 | 14.05263999 | 28 | $3.47 \times 10^{-9}$ |
| 8 | 15.40898658 | 29 | $3.82 \times 10^{-10}$ |
| 9 | 14.5637679 | 30 | $3.72 \times 10^{-11}$ |
| 10 | 12.00128216 | 31 | $3.19 \times 10^{-12}$ |
| 11 | 8.700584303 | 32 | $2.39 \times 10^{-13}$ |
| 12 | 5.58929941 | 33 | $1.54 \times 10^{-14}$ |
| 13 | 3.200105212 | 34 | $8.42 \times 10^{-16}$ |
| 14 | 1.640560267 | 35 | $3.84 \times 10^{-17}$ |
| 15 | 0.755903718 | 36 | $1.42 \times 10^{-18}$ |
| 16 | 0.313963174 | 37 | $4.07 \times 10^{-20}$ |
| 17 | 0.117823857 | 38 | $8.54 \times 10^{-22}$ |
| 18 | 0.04002034 | 39 | $1.16 \times 10^{-23}$ |
| 19 | 0.012318053 | 40 | $7.74 \times 10^{-26}$ |
| 20 | 0.00343814 | | |

**Table S2. Classification of the isolated ribozymes into five different types based on their ligase reactivities.**

| Isolated ligase ribozymes from *in vitro* evolution | | | | | |
|---|---|---|---|---|---|
| Cluster Peak sequences: in descending order of abundance (5′→3′) (only variable 40 nt region shown) | | No. of mutations from RS1 | Reaction catalyzed | Needs Template? | Classification |
| GAAUGCUGCCAACCGUGCGGGCUAAUUGGCAGACUGAGCU | RS1 | 0 | Ribozyme 3′ OH + Substrate 5′ AIP | Yes | Parent |
| ACGGGUGGGUAAUCUAGUGUCCGCGGAAUAGAACGAAACA | CS3 | 28 | 1) Ribozyme 3′ OH + Substrate 5′ PPP<br>2) Ribozyme 3′ OH + Substrate 5′AIP | Yes | Type I |
| GACAGCCGAGAAAUGAGUGGCCUAAAUGGGAGAAUGAGCU | CS1 | 13 | Substrate 2′ OH (3′ P) + Ribozyme 5′ PPP | No | Type IIa |
| GACUGCGCGUAUGAGUGGCGGCUAAAGAGGAGAAUGAGCG | CS2 | 16 | Substrate 2′OH (3′ P) + Ribozyme 5′ PPP | Yes | Type IIb |
| GGAUGGUGCGAACUGAGUGGGCUAAUUAGGAGAAUGAGCG | CS4 | 10 | Substrate 2′ OH (3′ P) + Ribozyme 5′ PPP | Yes | Type IIb |
| GGAGGGUGACAUCGUUGAGAGAGAAUGGGGAUAUUGAACU | CS5 | 16 | Substrate 2′ OH (3′ P) + Ribozyme 5′ PPP | Yes | Type IIb |
| GAAUCUGGCGAACGAUUAGUCCUAAUUGAGAAUUAUAGUU | CS8 | 18 | Substrate 2′ OH (3′ P) + Ribozyme 5′ PPP | Yes | Type IIb |
| AAGCUCUCGCCAGCAAAAGAACAGACCGUCGAGGAAACGG | CS6 | 28 | Ribozyme 3′ OH + Substrate 5′ AIP | Yes | Type III |
| CAAUGCUAUCCUCGGGGAACGAUUCUGCGGAAUCCGACAU | CS7 | 21 | Ribozyme 3′ OH + Substrate 5′ AIP | Yes | Type III |
| AAGUGAUGAAUUCCCUGCGGGCUACUUGUUAGAGCGGGCU | CS9 | 14 | Substrate Internal (11U) 2′ OH + Ribozyme 5′ PPP | Yes | Type IV |
| GUGUGUUACGAACCGUGGCGACUAAGCGGGAGGGUGAACU | CS10 | 14 | Substrate Internal (7G) 2′ OH + Ribozyme 5′ PPP | No | Type V |

**Table S3. Oligonucleotide sequences used in this work.** In the ribozyme sequences, variable nucleotides are highlighted in blue, the T7 promoter sequence is shown in purple, the hexauridine linker is shown in italics. 5′ and 3′ SHAPE cassettes are highlighted in green. Oligonucleotides were either purchased from Integrated DNA Technologies (IDT) or Chemgenes or generated enzymatically by *in vitro* transcription (IVT) of dsDNA templates (IDT). AIP-Substrate was generated by incubating the corresponding 5′ monophosphorylated RNA with EDC and 2-aminoimidazole (2AI) (see 'RNA preparation and substrate activation' in Materials and Methods).

| # | Oligo Name | Sequence (5′→3′) | Type | Source |
|---|---|---|---|---|
| 1.1 | Parent AI ligase (RS1) | GACUCACUGACACAGAUCCACUCACGGACAGCGGAAUGCUGCCAACCGUGCGGGCUAAUUGGCAGACUGAGCUCGCUGUCC*UUUUUU*GGCUAAGG | RNA | IVT |
| 1.2 | r0 DNA (Mutagenesis at 21% at each nucleotide position: 79% WT nucleotide and 7% of the other three) Note: The sequence is written according to IDT specifications | TAATACGACTCACTATAGACTCACTGACACAGATCCACTCACGGACAGCG(N1:07077907)(N2:79070707)(N2)(N3:07070779)(N1)(N4:07790707)(N3)(N1)(N4)(N4)(N2)(N2)(N4)(N4)(N1)(N3)(N1)(N4)(N1)(N1)(N1)(N4)(N3)(N2)(N2)(N3)(N3)(N1)(N1)(N4)(N2)(N1)(N2)(N4)(N3)(N1)(N2)(N1)(N4)(N3)CGCTGTCC*TTTTTT*GGCTAAGG | DNA | IDT |
| 2.1 | Template | GCGGUGGUCCUUAGCC | RNA | IDT |
| 2.2 | Modified Template | GCGGUGGUGGAAUCGC | RNA | IDT |
| 3.1 | PPPLigB | (5′-triphosphate)-ACCACCGCAUUCCGCA-(3′-BioTEG) | RNA | Chemgenes |
| 3.2 | PPPLig | (5′-triphosphate)-ACCACCGCAUUCCGCA | RNA | Chemgenes |
| 3.3 | PLig | 5′-monophosphate)-ACCACCGCAUUCCGCA | RNA | IDT |
| 3.4 | AIPLig | (5′-phosphoro-2-aminoimidazole)-ACCACCGCAUUCCGCA | RNA | Activation of PLig |
| 3.5 | PsLig | (5′-monophosphate)-ACCACCGC | RNA | IDT |
| 3.6 | AIPsLig | (5′-phosphoro-2-aminoimidazole)-ACCACCGC | RNA | Activation of PsLig |
| 3.7 | HOLigB | (5′-hydroxyl)-ACCACCGCAUUCCGCA-(3′-BioTEG) | RNA | IDT |
| 3.8 | HOLig | (5′-hydroxyl)-ACCACCGCAUUCCGCA | RNA | IDT |
| 3.9 | OHLig3′P | (5′-hydroxyl)-ACCACCGCAUUCCGCA-(3′-monophosphate) | RNA | IDT |
| 3.10 | OHLig3′P_dA16 | (5′-hydroxyl)-ACCACCGCAUUCCGCdA-(3′-monophosphate) | RNA | IDT |
| 3.11 | OHLig_dA16 | (5′-hydroxyl)-ACCACCGCAUUCCGCdA | RNA | IDT |
| 3.12 | OHLig_dC15 | (5′-hydroxyl)-ACCACCGCAUUCCGdCA | RNA | IDT |
| 3.13 | OHLig_dG14 | (5′-hydroxyl)-ACCACCGCAUUCCdGCA | RNA | IDT |
| 3.14 | OHLig_dC13 | (5′-hydroxyl)-ACCACCGCAUUCdCGCA | RNA | IDT |
| 3.15 | OHLig_dC12 | (5′-hydroxyl)-ACCACCGCAUUdCCGCA | RNA | IDT |
| 3.16 | OHLig_dU11 | (5′-hydroxyl)-ACCACCGCAUdUCCGCA | RNA | IDT |
| 3.17 | OHLig_dU10 | (5′-hydroxyl)-ACCACCGCAdUUCCGCA | RNA | IDT |
| 3.18 | OHLig_dA9 | (5′-hydroxyl)-ACCACCGCdAUUCCGCA | RNA | IDT |
| 3.19 | OHLig_dC8 | (5′-hydroxyl)-ACCACCGdCAUUCCGCA | RNA | IDT |
| 3.20 | OHLig_dG7 | (5′-hydroxyl)-ACCACCdGCAUUCCGCA | RNA | IDT |
| 3.21 | OHLig_dC6 | (5′-hydroxyl)-ACCACdCGCAUUCCGCA | RNA | IDT |
| 3.22 | OHLig_dC5 | (5′-hydroxyl)-ACCAdCCGCAUUCCGCA | RNA | IDT |

| # | Oligo Name | Sequence (5′→3′) | Type | Source |
|---|---|---|---|---|
| 3.23 | HOLigB_C16 | (5′-hydroxyl)-ACCACCGCAUUCCGCC-3'TEGBiot | RNA | IDT |
| 3.24 | HOLigB_G16 | (5′-hydroxyl)-ACCACCGCAUUCCGCG-3'TEGBiot | RNA | IDT |
| 3.25 | HOLigB_U16 | (5′-hydroxyl)-ACCACCGCAUUCCGCU-3'TEGBiot | RNA | IDT |
| 3.26 | HOLig_A11 | (5′-hydroxyl)-ACCACCGCAUACCGCA-3' | RNA | IDT |
| 3.27 | HOLig_C11 | (5′-hydroxyl)-ACCACCGCAUCCCGCA-3' | RNA | IDT |
| 3.28 | HOLig_G11 | (5′-hydroxyl)-ACCACCGCAUGCCGCA-3' | RNA | IDT |
| 3.29 | HOLig_A7 | (5′-hydroxyl)-ACCACCACAUUCCGCA-3' | RNA | IDT |
| 3.30 | HOLig_C7 | (5′-hydroxyl)-ACCACCCCAUUCCGCA-3' | RNA | IDT |
| 3.31 | HOLig_U7 | (5′-hydroxyl)-ACCACCUCAUUCCGCA-3' | RNA | IDT |
| 4.1 | PCR_Fwd_primer | TAATACGACTCACTATAGACTCACTGACAC | DNA | IDT |
| 4.2 | PCR_Rvs_primer | mCmCTTAGCCAAAAAAGGACAGCG | DNA | IDT |
| 4.3 | RT primer | GTGCGGAATGCGGTGGTCCTT | DNA | IDT |
| 4.4 | CSRvs_prim_mod | mGmGAATCGGAAAAAAGGACAGCG | DNA | IDT |
| 4.5 | CS_5′A_Fwd_primer | TAATACGACTCACTATTAACTCACTGACAC | DNA | IDT |
| 4.6 | SHAPE_RT_primer | (5′-FAM)-GAACCGGACCGAAGCCCG | DNA | IDT |
| 5.1 | CS1 | GACUCACUGACACAGAUCCACUCACGGACAGCGGACAGCCGAGAAAUGAGUGGCCUAAAUGGGAGAAUGAGCUCGCUGUCC*UUUUUU*GGCUAAGG | RNA | IVT |
| 5.2 | CS6 | GACUCACUGACACAGAUCCACUCACGGACAGCGAAGCUCUCGCCAGCAAAAGAACAGACCGUCGAGGAAACGGCGCUGUCC*UUUUUU*GGCUAAGG | RNA | IVT |
| 5.3 | CS6_5′t | GGACAGCGAAGCUCUCGCCAGCAAAAGAACAGACCGUCGAGGAAACGGCGCUGUCC*UUUUUU*GGCUAAGG | RNA | IVT |
| 5.4 | CS6_3′t | GACUCACUGACACAGAUCCACUCACGGACAGCGAAGCUCUCGCCAGCAAAAGAACAGACCGUCGAGGAAACGGCGCUGUCC | RNA | IVT |
| 5.5 | CS6_SHAPE | GGCCUUCGGGCCAAGACUCACUGACACAGAUCCACUCACGGACAGCGAAGCUCUCGCCAGCAAAAGAACAGACCGUCGAGGAAACGGCGCUGUCC*UUUUUU*GGCUAAGGUCGAUCCGGUUCGCCGGAUCCAAAUCGGGCUUCGGUCCGGUUC | RNA | IVT |
| 5.6 | CS7 | GACUCACUGACACAGAUCCACUCACGGACAGCGCAAUGCUAUCCUCGGGGAACGAUUCUGCGGAAUCCGACAUCGCUGUCC*UUUUUU*GGCUAAGG | RNA | IVT |
| 5.7 | CS7_5′t | GGACAGCGCAAUGCUAUCCUCGGGGAACGAUUCUGCGGAAUCCGACAUCGCUGUCC*UUUUUU*GGCUAAGG | RNA | IVT |
| 5.8 | CS7_3′t | GACUCACUGACACAGAUCCACUCACGGACAGCGCAAUGCUAUCCUCGGGGAACGAUUCUGCGGAAUCCGACAUCGCUGUCC | RNA | IVT |
| 5.9 | CS7_SHAPE | GGCCUUCGGGCCAAGACUCACUGACACAGAUCCACUCACGGACAGCGCAAUGCUAUCCUCGGGGAACGAUUCUGCGGAAUCCGACAUCGCUGUCC*UUUUUU*G | RNA | IVT |

| # | Oligo Name | Sequence (5′→3′) | Type | Source |
|---|---|---|---|---|
| | | GCUAAGGUCGAUCCGGUUCGCCGGAUCCAAAUCG GGCUUCGGUCCGGUUC | | |
| 5.10 | CS8 | GACUCACUGACACAGAUCCACUCACGGACAGCGG AAUCUGGCGAACGAUUAGUCCUAAUUGAGAAUUA UAGUUCGCUGUCC*UUUUUU*GGCUAAGG | RNA | IVT |
| 5.11 | CS8_5′t | GGACAGCGGAAUCUGGCGAACGAUUAGUCCUAAU UGAGAAUUAUAGUUCGCUGUCC*UUUUUU*GGCUAA GG | RNA | IVT |
| 5.12 | CS8_3′t | GACUCACUGACACAGAUCCACUCACGGACAGCGG AAUCUGGCGAACGAUUAGUCCUAAUUGAGAAUUA UAGUUCGCUGUCC | RNA | IVT |
| 5.13 | CS8_5′A | AACUCACUGACACAGAUCCACUCACGGACAGCGG AAUCUGGCGAACGAUUAGUCCUAAUUGAGAAUUA UAGUUCGCUGUCC*UUUUUU*GGCUAAGG | RNA | IVT |
| 5.14 | CS8_SHAPE | GGCCUUCGGGCCAAGACUCACUGACACAGAUCCA CUCACGGACAGCGGAAUCUGGCGAACGAUUAGUC CUAAUUGAGAAUUAUAGUUCGCUGUCC*UUUUUU*G GCUAAGGUCGAUCCGGUUCGCCGGAUCCAAAUCG GGCUUCGGUCCGGUUC | RNA | IVT |
| 5.15 | PPP-CS8_pc1 | (5′-triphosphate)- GACUCACUGACACAGAUCCACUCAC | RNA | IVT |
| 5.16 | P-CS8_pc1 | (5′-monophosphate)- GACUCACUGACACAGAUCCACUCAC | RNA | IDT |
| 5.17 | HO-CS8_pc1 | (5′-hydroxyl)- GACUCACUGACACAGAUCCACUCAC | RNA | IDT |
| 5.18 | P-CS8_pc2 | (5′-monophosphate)-GGACAGCGGAAUCUG GCGAACGAUUAGUCCUAAUUGAGAA | RNA | IDT |
| 5.19 | P-CS8_pc3 | (5′-monophosphate)-UUAUAGUUCGCUGUC CUUUUUUGGCUAAGG | RNA | IDT |
| 5.20 | CS8_splint_1 | CGCCAGATTCCGCTGTCCGTGAGTGGATCTGTGT CAG | DNA | IDT |
| 5.21 | CS8_splint_2 | CCAAAAAAGGACAGCGAACTATAATTCTCAATTA GGACTA | DNA | IDT |
| 5.22 | CS8_prim_mod | GACUCACUGACACAGAUCCACUCACGGACAGCGG AAUCUGGCGAACGAUUAGUCCUAAUUGAGAAUUA UAGUUCGCUGUCCUUUUUUCCGAUUCC | RNA | IVT |
| 5.23 | CS9 | GACUCACUGACACAGAUCCACUCACGGACAGCGA AGUGAUGAAUUCCCUGCGGGCUACUUGUUAGAGC GGGCUCGCUGUCC*UUUUUU*GGCUAAGG | RNA | IVT |
| 5.24 | CS9_5′t | GGACAGCGAAGUGAUGAAUUCCCUGCGGGCUACU UGUUAGAGCGGGCUCGCUGUCC*UUUUUU*GGCUAA GG | RNA | IVT |
| 5.25 | CS9_3′t | GGACAGCGAAGUGAUGAAUUCCCUGCGGGCUACU UGUUAGAGCGGGCUCGCUGUCC | RNA | IVT |
| 5.26 | CS9_5′A | AACUCACUGACACAGAUCCACUCACGGACAGCGA AGUGAUGAAUUCCCUGCGGGCUACUUGUUAGAGC GGGCUCGCUGUCC*UUUUUU*GGCUAAGG | RNA | IVT |
| 5.27 | PPP-CS9_pc1 | (5′-triphosphate)- GACUCACUGACACAGAUCCACUCAC | RNA | IVT |
| 5.28 | P-CS9_pc1 | (5′-monophosphate)- GACUCACUGACACAGAUCCACUCAC | RNA | IDT |
| 5.29 | HO-CS9_pc1 | (5′-hydroxyl)- GACUCACUGACACAGAUCCACUCAC | RNA | IDT |
| 5.30 | P-CS9_pc2 | (5′-monophosphate)-GGACAGCGAAGUGAU GAAUUCCCUGCGGGCUACUUGUUAG-3' | RNA | IDT |

| # | Oligo Name | Sequence (5′→3′) | Type | Source |
|---|---|---|---|---|
| 5.31 | P-CS9_pc3 | (5′-monophosphate)-AGCGGGCUCGCUGUCCUUUUUUGGCUAAGG-3' | RNA | IDT |
| 5.32 | CS9_splint_1 | TTCATCACTTCGCTGTCCGTGAGTGGATCTGTGTCAG | DNA | IDT |
| 5.33 | CS9_ splint_2 | CCAAAAAAGGACAGCGAGCCCGCTCTAACAAGTAGCCCG | DNA | IDT |
| 5.34 | CS9_prim_mod | GACUCACUGACACAGAUCCACUCACGGACAGCGAAGUGAUGAAUUCCCUGCGGGCUACUUGUUAGAGCGGGCUCGCUGUCCUUUUUUCCGAUUCC | RNA | IDT |
| 5.35 | CS10 | GACUCACUGACACAGAUCCACUCACGGACAGCGGUGUGUUACGAACCGUGGCGACUAAGCGGGAGGGUGAACUCGCUGUCC*UUUUUU*GGCUAAGG | RNA | IVT |
| 5.36 | CS10_5′t | GGACAGCGGUGUGUUACGAACCGUGGCGACUAAGCGGGAGGGUGAACUCGCUGUCC*UUUUUU*GGCUAAGG | RNA | IVT |
| 5.37 | CS10_3′t | GACUCACUGACACAGAUCCACUCACGGACAGCGGUGUGUUACGAACCGUGGCGACUAAGCGGGAGGGUGAACUCGCUGUCC | RNA | IVT |
| 5.38 | CS10_5′A | AACUCACUGACACAGAUCCACUCACGGACAGCGGUGUGUUACGAACCGUGGCGACUAAGCGGGAGGGUGAACUCGCUGUCC*UUUUUU*GGCUAAGG | RNA | IVT |
| 5.39 | PPP-CS10_pc1 | (5′-triphosphate)-GACUCACUGACACAGAUCCACUCAC | RNA | IVT |
| 5.40 | P-CS10_pc1 | (5′-monophosphate)-GACUCACUGACACAGAUCCACUCAC | RNA | IDT |
| 5.41 | HO-CS10_pc1 | (5′-hydroxyl)-GACUCACUGACACAGAUCCACUCAC | RNA | IDT |
| 5.42 | P-CS10_pc2 | (5′-monophosphate)-GGACAGCGGUGUGUUACGAACCGUGGCGACUAAGCGGGAG | RNA | IDT |
| 5.43 | P-CS10_pc3 | (5′-monophosphate)-GGUGAACUCGCUGUCCUUUUUUGGCUAAGG | RNA | IDT |
| 5.44 | CS10_ splint_1 | CGTAACACAACCGCTGTCCGTGAGTGGATCTGTGTCAG | DNA | IDT |
| 5.45 | CS10_ splint_2 | CCAAAAAAGGACAGCGAGTTCACCCTCCCGCTTAGTCGCC | DNA | IDT |

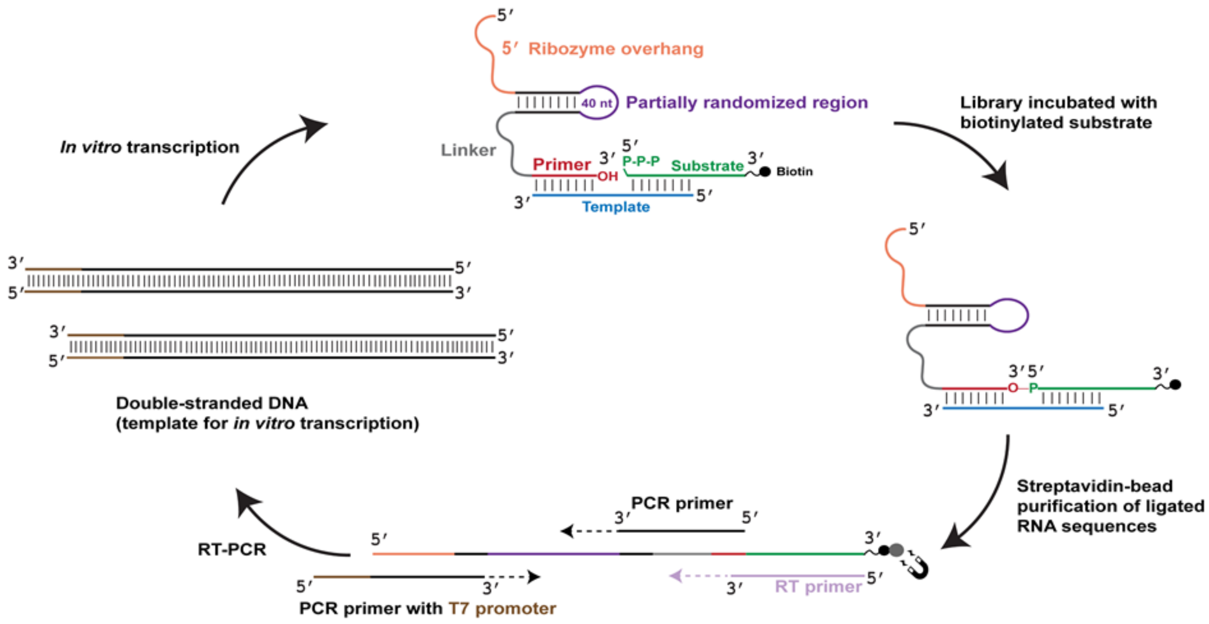


**Fig. S1. Selection protocol used to isolate ligase ribozymes.** An RNA library, containing a partially randomized region (depicted in dark purple) derived from an AIP-ligase,[1] was challenged with an RNA substrate (depicted in green), containing a 5′-triphosphate group and a 3′-TEG-biotin group (depicted in black) in the presence of an external RNA template (depicted in blue). Ligated sequences were purified by capturing them on streptavidin-coated magnetic beads (depicted as a gray circle) and reverse transcribed using a primer (RT primer; depicted in light purple) that is complementary to the entire substrate sequence. The RT primer also has the potential to bind directly to the library sequences by forming four base-pairs with the 3′ end of its 'primer' region (depicted in red). The cDNA was PCR-amplified, with the T7 promoter sequence (depicted in brown) added to the dsDNA sequence during PCR. This dsDNA was transcribed to generate the library for subsequent rounds of selection.

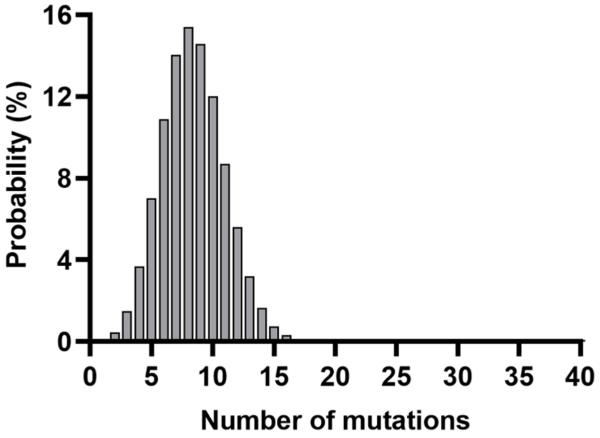


**Fig. S2.** Distribution of sequence variants of the ancestral AIP-ligase, RS1, in the starting RNA library that was doped at 21%.

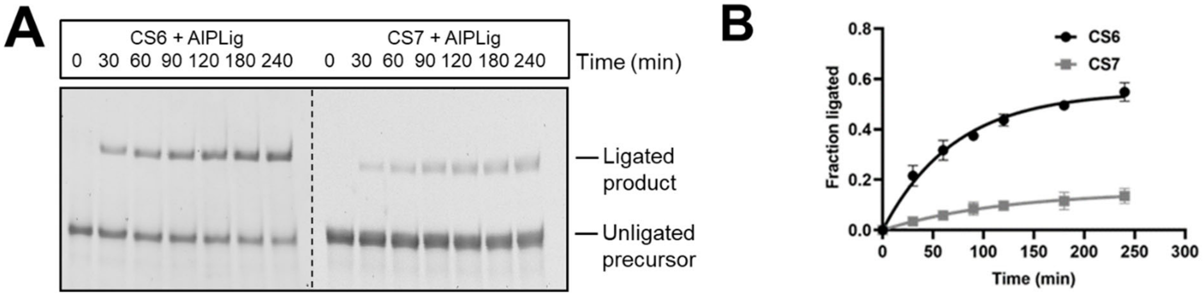


**Fig. S3. Kinetics of CS6- and CS7-catalyzed ligation with AIPLig. A.** Time course of ligation catalyzed by CS6 and CS7. CS6 is a more efficient ligase than CS7. **B.** Kinetic plot for CS6- and CS7-catalyzed ligation. CS6 and CS7 showed ligation yields of 60% and 10%, respectively, after 4 h, and CS6 showed faster kinetics. Ligation reactions contained 1 μM ribozyme, 1.2 μM RNA template, and 2 μM AIPLig in 100 mM Tris-HCl (pH 8.0), 300 mM NaCl, and 10 mM $MgCl_2$. Experiments were performed in triplicate; error bars indicate standard deviation.

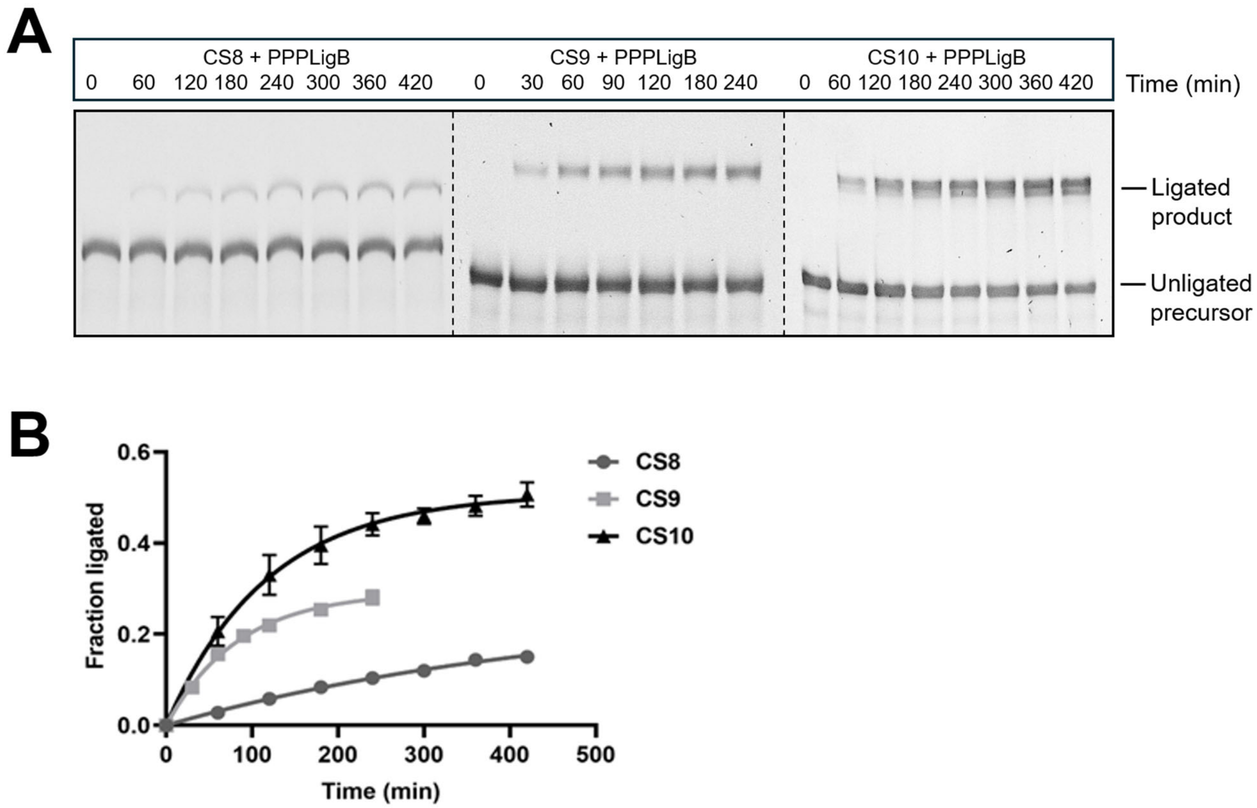


**Fig. S4. Kinetics of RNA ligation with PPPLigB catalyzed by CS8, CS9, and CS10. A.** Time course of ligation catalyzed by CS8, CS9, and CS10. **B.** Kinetic plots for ligation catalyzed by CS8, CS9, and CS10. CS9, and CS10 exhibited comparable rates, with CS10 producing more ligated product. CS8 was less efficient, generating 10% ligated product after 4 h. Ligation reactions contained 1 µM ribozyme, 1.2 µM RNA template, and 2 µM PPPLigB in 100 mM Tris-HCl (pH 8.0), 300 mM NaCl, and 100 mM $MgCl_2$. Experiments were performed in triplicate; error bars indicate standard deviation.

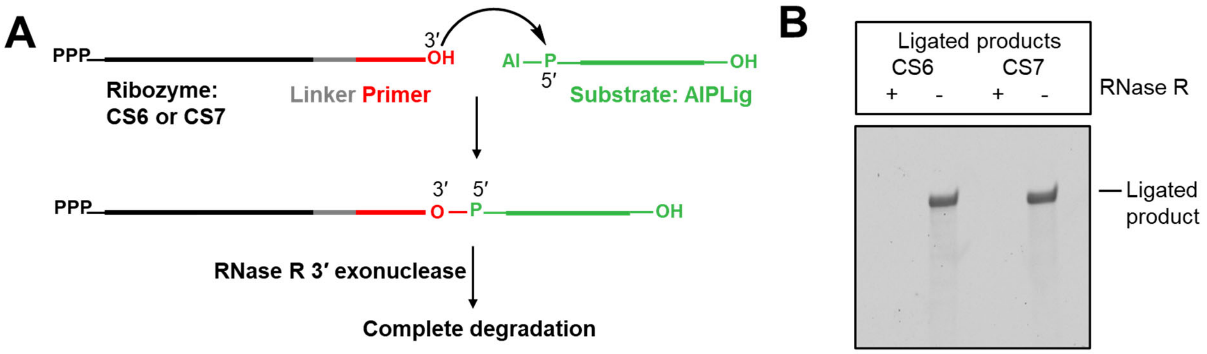


**Fig. S5. Ligation creates a 3′-5′ phosphodiester bond between the substrate and ribozymes, CS6 and CS7.** **A.** Schematic of the 3′ exonuclease assay to probe the regiochemistry of the phosphodiester bond between the ribozyme and substrate after ligation. **B.** The purified ligated product from the reaction between CS6 or CS7 and AIPLig was completely digested by the 3′→5′ exonuclease, RNase R, indicating that ligation by CS6 and CS7 generates a canonical 3′-5′ bond. Exonuclease digestion was carried out according to vendor specifications (see ‘Confirmation of 3′-5′ phosphodiester bonds in the ligated products generated by CS6 and CS7’ in ‘Materials and Methods’).

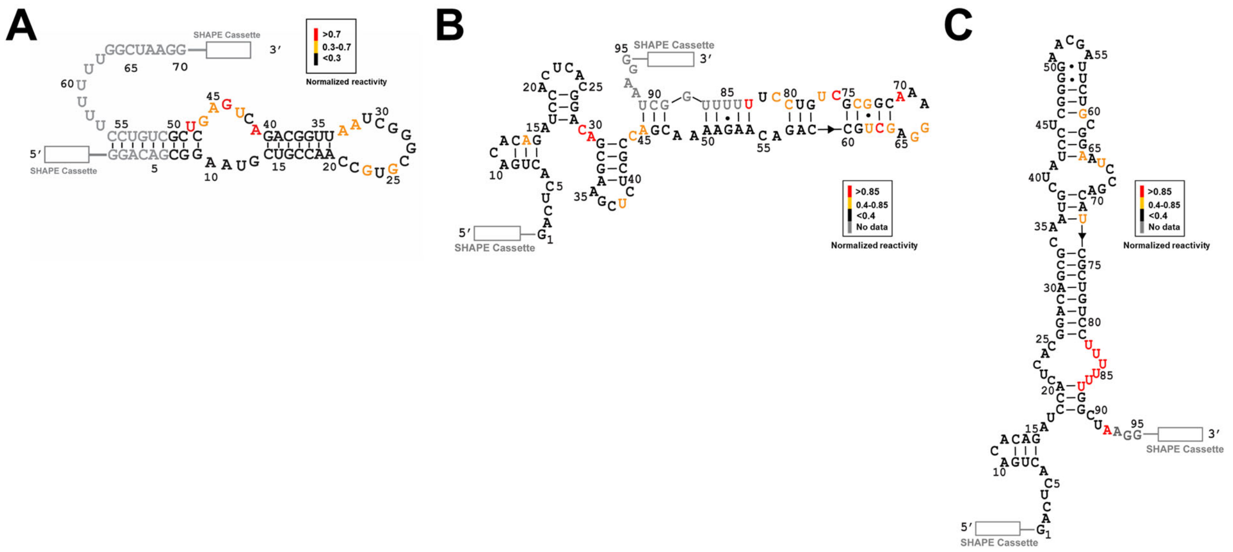


**Fig. S6. SHAPE analysis of RS1, CS6, and CS7.** SHAPE-derived secondary structures of (A) RS1[1], (B**)** CS6, and (C) CS7. All structures were determined by the RNAstructure program using reactivity constraints obtained from SHAPE experiments.[2] 5′ and 3′ SHAPE cassettes are denoted by white rectangles. Nucleotides for which no data were obtained are shown in gray. Nucleotides are colored in red, orange, and black according to their normalized SHAPE reactivities, as shown in the reactivity legend. Sequences used in SHAPE experiments can be found in Table S3. All three ribozymes catalyze ligation using the same chemical pathway but have divergent sequences and adopt distinct secondary structures.

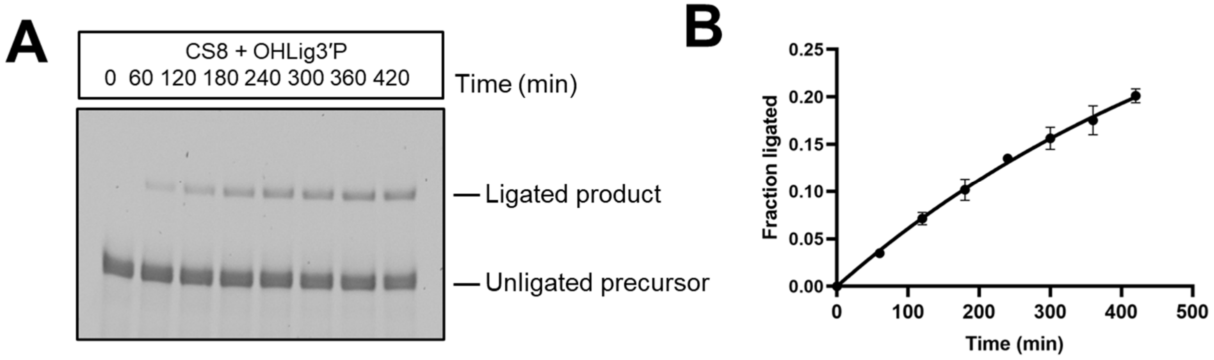


**Fig. S7. Kinetics of CS8-catalyzed ligation with OHLig3′P. A.** Time course of ligation catalyzed by CS8. **B.** Kinetic plot for ligation catalyzed by CS8. Ligation reactions contained 1 μM ribozyme, 1.2 μM RNA template, and 2 μM RNA substrate in 100 mM Tris-HCl (pH 8.0), 300 mM NaCl, and 100 mM $MgCl_2$. Experiments were performed in triplicate; error bars indicate standard deviation.

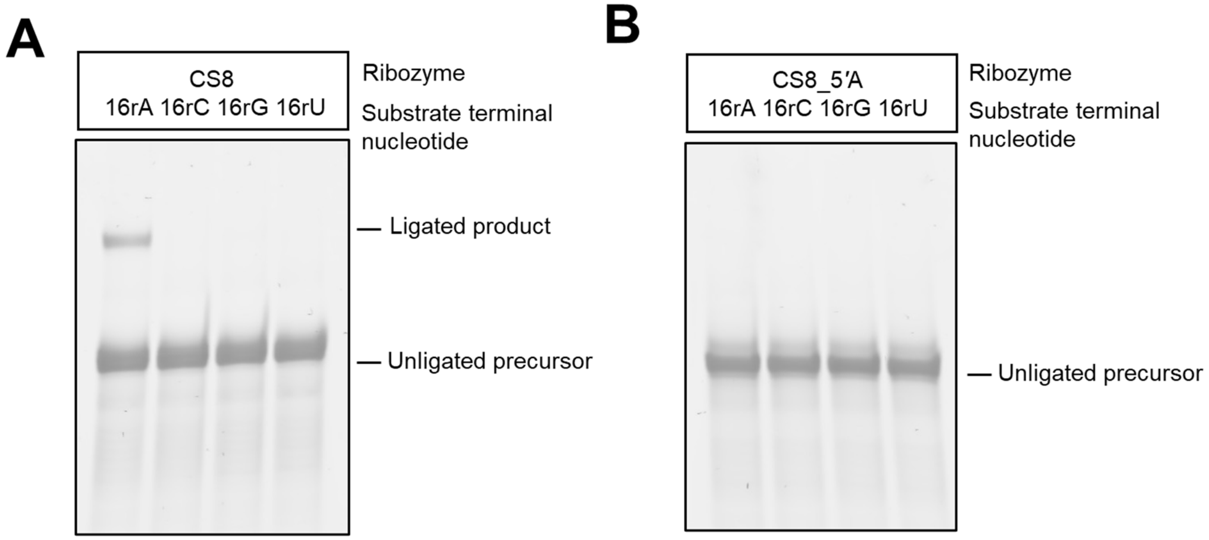


**Fig. S8. Nucleotide requirements at the ligation junction for CS8-catalyzed RNA ligation.** CS8-catalyzed ligation requires a (A) 3′ terminal A on the substrate and a (B) 5′ terminal G on the ribozyme.

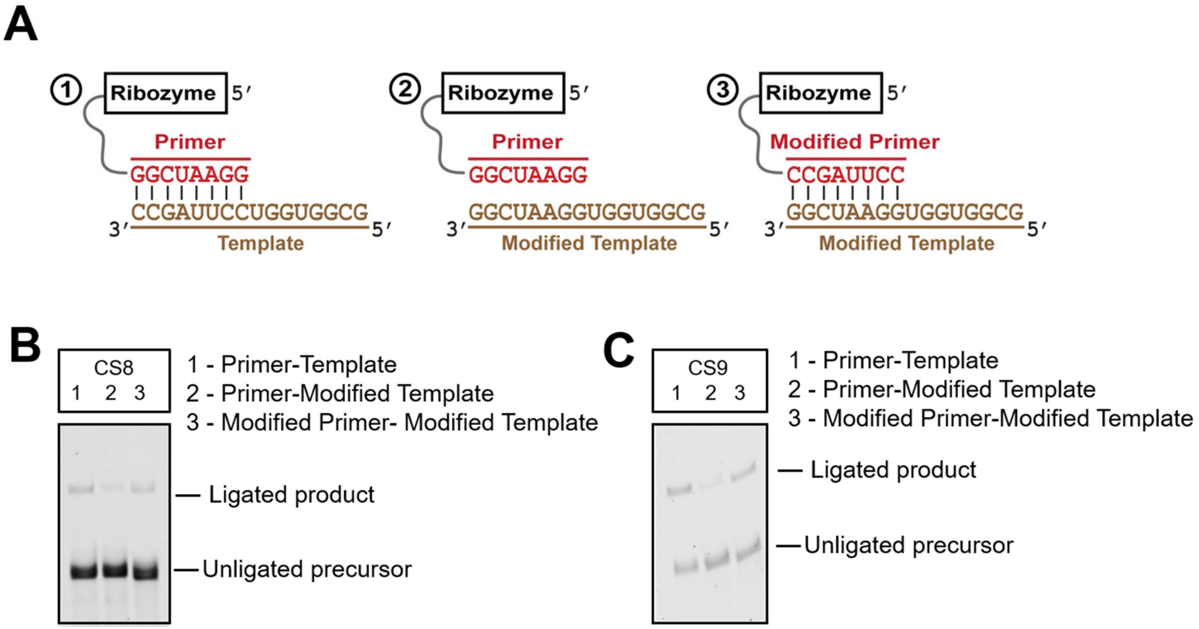


**Fig. S9. Template-assisted RNA ligation by CS8 and CS9. A.** Schematic for the compensatory mutational rescue of ligase activity in CS8 and CS9. Base-pairing interactions between the template and the 3′-primer sequence of (B) CS8 and (C) CS9 are important for ligation. Disrupting this interaction by mutations to the template significantly reduces ligation; however, compensatory mutations in the 3′-primer sequence of the ribozymes rescue ligation. Ligation reactions contained 1 μM ribozyme, 1.2 μM RNA template, and 2 μM substrate (OHLig for CS9 and OHLigB for CS8) in 100 mM Tris-HCl (pH 8.0), 300 mM NaCl, and 100 mM $MgCl_2$. Ligations were assayed at 3 h.

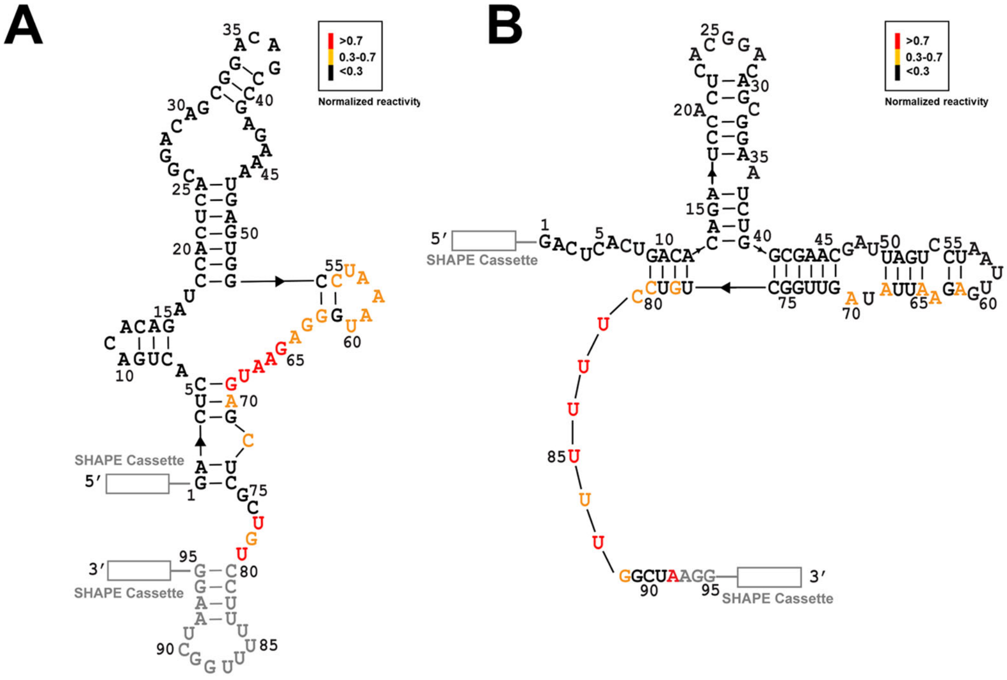


**Fig. S10. SHAPE analysis of CS1 and CS8. A.** Secondary structure of CS1.[3] **B.** Secondary structure of CS8. Structures were determined by the RNAstructure program using reactivity constraints obtained from SHAPE experiments.[3] 5′ and 3′ SHAPE cassettes are denoted by white rectangles. Nucleotides for which no data were obtained are shown in gray. Nucleotides are colored in red, orange, and black according to their normalized SHAPE reactivities, as shown in the reactivity legend. Sequences used in SHAPE experiments can be found in Table S3. Although both ligases exhibit the same chemical reactivity, they are separated by 22 mutations and adopt distinct secondary structures.

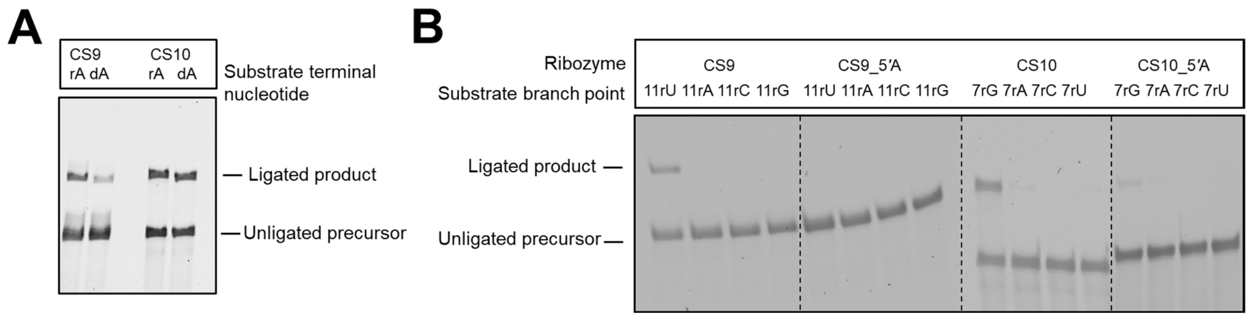


**Fig. S11. CS9- and CS10-catalyzed ligation reactions involve conserved internal nucleotides. A.** CS9- and CS10-catalyzed ligation is preserved when the 3′ terminal nucleotide (A16) of the substrate is substituted by deoxyadenine, indicating that A16 does not harbour the nucleophile for this reaction. **B.** The 5′ terminal guanine (the residue that contains the 5′-triphosphate group) of CS9 and an internal uracil (U11 that contains the nucleophilic 2′-hydroxyl group) of the substrate are essential for ligation by CS9. CS10 withstands a G1A mutation with marked reduction in ligation, but both 'wild-type' and mutant ribozymes require internal guanines at the 7$^{th}$ position (G7) of the substrate. Ligation reactions contained 1 µM ribozyme, 1.2 µM RNA template, and 2 µM OHLig in 100 mM Tris-HCl (pH 8.0), 300 mM NaCl, and 100 mM $MgCl_2$. Ligations were assayed at 3 h.
